# Small protein 26 interacts and enhances glutamine synthetase activity in *Methanosarcina mazei*

**DOI:** 10.1101/2020.05.06.080432

**Authors:** Miriam Gutt, Britta Jordan, Katrin Weidenbach, Mirja Gudzuhn, Claudia Kiessling, Liam Cassidy, Andreas Helbig, Andreas Tholey, Dennis Pyper, Harald Schwalbe, Ruth A. Schmitz

**Author notes:** joined first, equal contribution (Miriam Gutt, Britta Jordan). Corresponding author:* Ruth A. Schmitz, Institut für Allgemeine Mikrobiologie, Christian-Albrechts-Universität zu Kiel, Am Botanischen Garten 1-9, 24118 Kiel, Germany.

## Abstract

Small ORFs (sORF) encoded small proteins have been overlooked for a long time due to challenges in prediction and distinguishing between coding and non-coding predicted sORFs and in their biochemical detection and characterization. We report on the first biochemical and functional characterization of a small protein (sP26) in the archaeal model organism *Methanosarcina mazei*, comprising 23 amino acids. The corresponding encoding leaderless mRNA (spRNA26) is highly conserved within numerous Methanosarcina strains on the amino acid as well as on nucleotide level strongly arguing for a cellular function of the small protein. spRNA26 is significantly enhanced under nitrogen limitation, but also under oxygen and salt stress conditions. His-tagged sP26 was heterologously expressed and purified by fractionated ammonium sulfate precipitation, affinity chromatography and size exclusion centrifugation. Using independent biochemical approaches (pull-down by affinity chromatography followed by MS analysis, revers pull-down, microscale thermophoresis and size exclusion chromatography) we observed that sP26 interacts and forms complexes with *M. mazei* glutamine synthetase (GlnA_1_) with high affinity (app. KD = 45 +/− 14 µM). Upon interaction with sP26, GlnA_1_ activity was significantly stimulated independently and in addition to the known activation by the metabolite 2-oxoglutarate. Besides strong interaction of sP26 with the PII-like protein GlnK_1_ was demonstrated (KD= 1.4 µM +/− 0.9 µM). On the basis of these findings, we hypothesize that in addition to 2-oxoglutarate, sP26 activates GlnA_1_ activity under nitrogen limitation most likely by stabilizing the dodecameric structure of GlnA_1_.

## INTRODUCTION

Modern genomics and transcriptomics technologies combined with systematic genome-wide approaches have uncovered an unexpected genome complexity in prokaryotes. Besides genes encoding larger proteins and genes of non-coding RNAs (ncRNAs), global approaches have over the past decade discovered a wealth of hidden small genes containing short open reading frames (sORFs) in many prokaryotic genomes (Hobbs et al. (2011), Ramamurthi and Storz (2014), recently reviewed in Orr et al. (2020)). These sORFs often encode proteins smaller than 50 amino acids (aa) in length, and have been typically missed in genome annotations by automated gene predictions due to too strict assumptions and traditionally considering only standard proteins in the automated gene annotation tools (Basrai et al. (1997), Miravet-Verde et al. (2019)). This led to paying no attention to the existence of an additional layer of complexity represented by the small proteins. Besides, those small proteins have been difficult to detect biochemically due to technical limitation. Though, nowadays new technologies are emerging, which enable their global profiling in genome-wide approaches, and systematic approaches for global identification are used e.g. by sophisticated bioinformatics predictions, peptidomics, ribosome profiling and combinations of those (e.g. Cassidy et al. (2019), Cassidy et al. (2016), Lago et al. (2017), Saghatelian and Couso (2015), Chu et al. (2015), Cardon et al. (2020), Ma et al. (2014), Ingolia (2014), Meydan et al. (2019), Weaver et al. (2019), Miravet-Verde et al. (2019)). Due to their small size the small proteins are frequently predicted to modify the activity of larger proteins or complexes via physical interactions or interact with the membranes. However, information regarding specific physiological role(s) of verified small proteins is lacking for the majority of confirmed small proteins. Only a fraction of small proteins experimentally identified in diverse prokaryotes have been functionally characterized. This characterization demonstrated that they can play important roles in different functional scenarios and have a broad range of function from cell division, signal transduction, modeling membrane protein recruitment or (membrane) protein activity to modulation or being part of a larger mostly membrane integrated protein complex (reviewed by Storz et al. (2014), Duval and Cossart (2017), Orr et al. (2020)). In archaea a few small proteins have been reported, the majority of which is regulated in response to a specific stress. However, for most of them verified functional analysis is scarce or missing (Prasse et al. (2015), Cassidy et al. (2016), Cassidy et al. (2018), Nagel et al. (2019), Kubatova et al. (2019), Jevtic et al. (2019))

*Methanosarcina mazei* strain Gö1 belongs to the methylotrophic methanogens of the order *Methanosarcinales*, which have the most versatile substrate spectrum within the methanogenic archaea and significantly contribute to the production of the greenhouse gas (Ferry (1999)). *M. mazei* is able to fix molecular nitrogen under nitrogen limitation. The regulation of the nitrogen metabolism particularly of the nitrogen fixation as well as the glutamine synthetase is well studied on the transcriptional and post-transcriptional level (Jager et al. (2009), Prasse et al. (2017), Buddeweg et al. (2018), Weidenbach et al. (2008), Weidenbach et al. (2010), Weidenbach et al. (2014)). Overall regulation occurs in response to the nitrogen (N) availability, where N limitation is in general perceived internally by sensing the internal 2-oxoglutarate pool, which increases under N limitation due to reduced consumption by the ammonium-dependent glutamate dehydrogenase and increasing glutamine synthetase / glutamate synthase pathway (GS/GOGAT) for glutamate synthesis (Ehlers et al. (2005b)). We recently showed that post-transcriptional regulation by small RNA_154_, originally identified in a global RNAseq approach (Jager et al. (2009)), plays a central role in N regulation of several components of the N cycle including glutamine synthetase (Prasse et al. (2017)). Besides, RNAseq and term-seq approaches not only identified high numbers of small non-coding RNAs but also numerous small mRNAs in *M. mazei* containing putative sORFs (Dar et al. (2016), Jager et al. (2009) as well as unpublished RNAseq data sets). Aiming to elucidate whether the predicted sORFs identified are translated *in vivo*, the full cytosolic proteome of *M. mazei* was examined using reversed phase LC-MS MS approaches (bottom up and top down strategies) as described in Prasse et al. (2015), Cassidy et al. (2019), Cassidy et al. (2016)). In these comprehensive studies overall 40 small proteins were detected and experimentally validated with high or mid confidence (Cassidy et al. (2019), Cassidy et al. (2016)). In this present study the tentatively identified small protein 26 (sP26), which was identified with low confidence in our previous study, was selected aiming to identify its physiological role, due to the fact that the respective encoding spRNA26 is upregulated under N starvation (Jager et al. (2009)). Moreover, the genomic location of spRNA26 down-stream of the operon encoding the acetyl-CoA-decarbonylase/synthase (ACS) complex (Jager et al. (2009)), suggested ACS, which itself is up-regulated in response to the N starvation on the post-transcriptional level (Buddeweg et al. (2018)), as an attractive potential target for sP26. Very recently NMR spectroscopy analysis demonstrated that sP26 is unstructured, however bioinformatic tools predict that unstructured sP26 can potentially fold into a structure upon complex formation with a target (Kubatova et al. (2020)). Here we demonstrate by independent biochemical approaches that sP26 interacts with glutamine synthetase (GlnA_1_), the central component of the nitrogen cycle in *M. mazei*, and stimulates its activity.

## MATERIALS AND METHODS

### Strains and plasmids

All plasmids used have been constructed as follows, confirmed by sequencing and are summarized in table 1. pRS1242 was generated for overproducing sP26 N-terminally fused to a (His)_6_-tag (His_6_-sP26) in *M. mazei*. A DNA fragment including the respective gene was commercially synthesized (Eurofins Scientific, Nantes, Luxemburg) as follows: The gene was placed under the control of the promoter and the ribosome binding site of *mcr*B (Gernhardt et al. (1990)), followed by an additional ATG as new translation start and the sORF26 with an N-terminal His_6_-Tag. Downstream of the sORF26 sequence ending with the native TAA the methanoarchaeal transcriptional terminator (TTTT) was added. Further a flanking *Sac*I restriction site at the 5’ and a *Kpn*I restriction site at the 3’ end were added resulting in a 379 bp fragment. The synthesized DNA fragment was delivered inserted into pEX (Eurofins Scientific, Nantes, Luxemburg) the corresponding plasmid designated pRS1209 (see Supp. Fig. S2). Using the restriction endonucleases *Sac*I and *Kpn*I (NEB, Schwalbach, Germany) the respective 373 bp fragment was generated and ligated into the *Sac*I and *Kpn*I restricted pWM321 vector (Metcalf et al. (1997)) resulting in plasmid pRS1242 (see Supp. Fig. S2).

**Table 1.** Strains and plasmids used.

| Strain or plasmid | Genotype or description | Source of reference |
| --- | --- | --- |
| **Strains** |  |  |
| *Methanosarcina mazei* strain Gö1 | wild type; DSM No. 3647 | DSMZ, Braunschweig, Germany |
| *Escherichia coli* BL21 (DE3) | general expression strain | Invitrogen, Carlsbad, USA |
| *Escherichia coli* BL21-CodonPlus®-RIL | general expression strain containing the pRIL plasmid (ileW, leuY, proL) | Stratagene, La Jolla, USA |
| **Plasmids** |  |  |
| pET28a | general cloning vector providing N-terminal His<sub>6</sub>-tag | Merck KGaA, Darmstadt, Germany |
| pET-SUMO | general cloning vector providing N-terminal SUMO-tag | Invitrogen, Carlsbad, USA |
| pCR®II –TOPO® vector | general cloning vector | Life Technologies, Darmstadt, Germany |
| pRS196 | MM0964 (*glnA<sub>1</sub>*) under the control of T7 in pET28a | Ehlers et al. (2005b) |
| pRS203 | MM0732 (*glnK<sub>1</sub>*) under the control of T7 in pET28a | Ehlers et al. (2005b) |
| pRS375 | pRS196 with exchanged His<sub>6</sub>-Tag by Strep-tag | this work |
| pRS1209 | His<sub>6</sub>-sORF26 under the control of *pmcrB* in pEX | Eurofins Scientific, Nantes, Luxemburg |
| pRS1228 | SUMO-sORF26 under the control of T7 in pET-SUMO | this work |
| pRS1229 | SUMO-His<sub>6</sub>-sORF26 under the control of T7 in pET-SUMO | this work |
| pRS1242 | His<sub>6</sub>-sORF26 under the control of *pmcrB* in pWM321 | this work |
| pRS1244 | sORF26 in pCR®II-TOPO® | this work |
| pRS1245 | His<sub>6</sub>-sORF26 under the control of T7 in pET28a | this work |

For production of sORF26 fused to a SUMO-Tag pRS1209 as template, the primers sORF26_1 His-for (5’-ATGCATCATCATCATCATCACGTGCCTGTGATGAAAAATCTG-3’) and sORF26_1 rev (5’-AAAAAATTAAGGATAATTTCCGTGCCTC-3’) were used in a PCR to generate a 100 bp fragment encoding sORF26 with an N-terminal His_6_-Tag. This fragment was TA-cloned into the pET-SUMO (Invitrogen, Carlsbad, USA) resulting in pRS1229 (see Supp. Fig. 2). The respective resulting fusion protein is an N-terminal His_6_-SUMO-Tag fused to His_6_-sP26 (16.3 kDa).

pRS1228 was constructed by amplifying sORF26 from pRS1209 using primers sORF26_1 rev (AAAAAATTAAGGATAATTTCCGTGCCTC) and sORF26_1 for (GTGCCTGTGATGAAAAATCTGGCTG) and TA-cloning the respective fragment into pET-SUMO (Invitrogen, Carlsbad, USA) vector. In contrast to pRS1229, the respective construct pRS228 provides no additional His_6_-tag (see Supp. Fig. S2).

pRS1245 for expression of His_6_-sORF26 in *E. coli* was derived from plasmid pRS1209 as follows. The sequence for sORF26 was PCR amplified using pRS1209 as template and the primers sORF26_1_for_*Nde*I (5’-<u>CATATG</u>GTGCCTGTGATGAAAAATCTGGCTG-3’) and sORF26_1_rev_*Nde*I (5’-CATATGGAAAAAATTAGGATAATTTCCGTGCCTC-3’) which add flanking *Nde*I restriction sites (underlined). The resulting 91 bp product was TA-cloned into the vector pCR®II-TOPO® (Life Technologies, Darmstadt, Germany) creating the plasmid pRS1244. After *Nde*I restriction of pRS1244 (NEB, Schwalbach, Germany) the generated 84 bp fragment was cloned into the *Nde*I linearized vector pET28a(+) (Merck KGaA, Darmstadt, Germany) resulting in pRS1245 where the N-terminal vector derived His_6_-Tag followed by a thrombin site is fused to sORF26 (see supplemental Fig. S2). The corresponding small protein has a calculated molecular mass of 5.45 kDa.

Plasmid pRS375 was constructed by amplifying *glnA*_*1*_ from pRS196 (Ehlers et al. (2005b)) using a forward primer replacing the N-terminal (His)_6_-tag by a Streptag (Mm GlnA H/S for 5’-CCATGGGCTGGAGCCACCCGCAGTTCGAAAAAAGCAGCGGCCTGGTGCCGCGC-3’) and adding a *Nco*I restriction site, and the reverse primer (Mm GlnA H/S rev 5’-CCGCCGCGCCCGCCGCCCGCCGCAAGCTTGTCGACGGAGCTCGAATTCGGATCC-3’) adding a *Bam*HI restriction site. The obtained 1.4 kbp PCR product was cloned into pRS196 using *Nco*I and *Bam*HI restriction sites. The resulting plasmid was designated pRS375.

For expression in *E. coli* all plasmids were transformed into *E. coli* BL21(DE3) (Invitrogen, Carlsbad, USA) or *E. coli* BL21 pRIL (Stratagene, La Jolla, USA) using the method of Inoue et al. (1990). For overproduction in *M. mazei* the plasmid pRS1242 was transformed into *M. mazei* DSM3647 by liposome-mediated transformation as described by Ehlers et al. (2005a).

### Growth analysis of *M. mazei*

All *M. mazei* strains were grown at 37 °C in 50 ml or 1 L enclosed serum bottles under anaerobic conditions, with the gaseous phase containing N_2_ and CO_2_ (80/20) (Air Liquide, Paris, France). The minimal medium was supplemented with 150 mM methanol and 40 mM acetate as carbon and energy sources, as described in Veit et al. (2006). For growth under N limitation the N_2_ from the gas atmosphere served as the sole N source and for N sufficiency the medium was supplemented with 10 mM NH_4_Cl. For different stress conditions the cultures were grown under various conditions, at 30 °C, at 42 °C, under oxygen stress (added 20 ml sterile air at a turbidity of 0.25), and under salt stress (additional 500 mM NaCl). Growth was generally monitored by determining the turbidity of the cultures at 600 nm.

### RNA isolation, northern and RNAseq analysis

RNA isolation was performed using Rotizol reagent (Carl Roth GmbH, Karlsruhe, Germany) following the manufacturer’s protocol. Subsequently a DNaseI treatment and phenol-chloroform-isoamylalcohol precipitation as described in Nickel et al. (2013). Northern Blot analysis was performed as described by Jager et al. (2009). RNAs were detected using 5’-^32^P labeled ssDNA oligo probes against the sORF26 (5’GCCTGTGATGAAAAATCTGGCTG3’) and against the 5S rRNA (5’CGCACACTTCAGTACAGTAAGGAA3’). The pGEM™ Conventional DNA Digest Marker PR-G1741 (Promega™, Fitchburg, USA) was used as size standard and the bands were quantified by the AIDA Software (Raytest, Staubenrath, Germany).

The Illumina sequencing data were obtained from *M. mazei* cultures grown under N starvation which were harvested in the exponential growth phase. RNA isolation was performed as described above and RNAsequencing was performed as described by Prasse et al. (2017).

### Purification of SUMO-sP26 and co-elution analysis

His_6_-SUMO-His_6_-sP26 was purified from 1 L culture of *E. coli* BL21 (DE3) carrying the plasmid pRS1229 and grown in LB at 37 °C. At a turbidity of 0.6 at 600 nm the medium was supplemented with 100 µM IPTG and further incubated for 2 h at 37°C with rigorous shaking. After harvesting the cells (4.000 x *g* at 4 °C for 30 min) and resuspending in buffer A (50 mM NaH_2_PO_4_, 300 mM NaCl, pH 8). The cells were disrupted using a French pressure cell two times (at 4.135 x 10^6^ N/m^2^) followed by centrifugation at 13.865 x *g* at 4 °C for 30 min. The cytosolic supernatant was incubated and stirred with 1 ml Ni-NTA agarose (Qiagen, Hilden, Germany) at 4 °C for 1 h. After pouring the slurry in an empty column, it was washed twice with 8 ml buffer A containing 20 mM imidazole. The SUMO-His_6_-sP26 protein was eluted with 5 × 1 ml buffer A containing 100 mM, 250 mM and 500 mM imidazole. Elution fractions containing the SUMO-His_6_-sP26 protein (250 mM imidazole) were combined and dialyzed overnight against buffer A. For co-elution experiments 1 mg of purified SUMO-His_6_-sP26 protein was incubated with 500 µl *M. mazei* crude cell extract (27.5 mg) and immobilized to 0.5 ml Ni-NTA agarose for 1 h at 4°C. All non-binding proteins were washed off with 5 x 1 ml buffer A and the SUMO-His_6_-sP26 and potentially interacting proteins eluted with 4 x 1 ml buffer A supplemented with 250 mM imidazole. The elution fractions were analyzed with Coomassie and silver stained SDS-PAGE and Tris-Tricine-PAGE (see Gel electrophoretic separation). The respective *M. mazei* crude cell extract was generated as follows: 1 L *M. mazei* culture grown exponentially under N limitation was harvested at 4.000 *x g* and 4 °C for 30 min and the cell pellet resuspended in 500 µl 50 mM Tris/HCl pH 6.8 supplemented with DNaseI (Thermo Fisher Scientific™, Waltham, USA). Cells were twice disrupted by GinoGrinder (SPEX CertiPrep, Metuchen, USA) on ice at 1,300 strokes for 3 min each, followed by centrifugation at 13.865 x *g* and 4 °C for 30 min to separate cell debris.

### Identification of co-eluting proteins by LC-MSMS analysis

Coomassie blue (R-250) stained gel bands were excised and destained. The individual bands were reduced with dithiothreitol (10 mM, 56°C, 1 hr), and subsequently alkylated in the presence of iodoacetamide (50 mM, room temperature, 30 min). Enzymatic digestion of was performed overnight at 37°C by the addition of sequencing grade trypsin (Promega, Wisconsin) (100 ng per sample) in 100 µl of ammonium bicarbonate (ABC) buffer (100 mM pH 7.4). The peptides were extracted via one wash with 60% acetonitrile with 0.1% trifluoroacetic acid, and a second wash with 100% acetonitrile. The peptides were then dried via vacuum evaporation prior to LC-MS analysis. For LC-MS analysis the samples were suspended in UHPLC loading buffer (3% acetonitrile + 0.1% trifluoroacetic acid).

Chromatographic separation was performed on a Dionex U3000 UHPLC system (Thermo Fisher Scientific, Darmstadt, Germany) equipped with an Acclaim PepMap 100 column (3 μm particle size, 75 μm × 150 mm) and µ-pre-column (300 μm × 5 mm) coupled online to LTQ Orbitrap Velos mass spectrometer (Thermo Fisher Scientific). The eluents used were; eluent A: 0.05% formic acid (FA), eluent B: 80% ACN + 0.05% FA. The separation was performed over a programmed 90-min run. Initial chromatographic conditions were 5% B for 5 minutes followed by an increase to 10% B over 1 min, subsequently a linear gradient from 10% to 40% B over 54 min, a 5-min increase to 95% B, and 10 min at 95% B. Following this, an inter-run equilibration of the column was achieved by 15 min at 5% B. A constant flow rate of 300 nl/min was used and 8 μl of sample was injected per run. Data acquisition on the LTQ Orbitrap Velos mass spectrometer utilized CID activation (NCE 35). A full scan MS acquisition was performed (resolution 60,000) scan range 300-1500 m/z, maximum IT 100 ms. Subsequent data dependent MS/MS (resolution 7,500, minimum intensity 500,), of the top 10 most intense ions, single charged and undetermined charged state ions were excluded, dynamic exclusion was enabled (90 sec duration, repeat count 2 in 30 sec); internal lockmass was enabled on 445.12003 m/z.

MS data files were searched against a FASTA-database containing the full *M. mazei* proteome (accessed from UniProt 2016.03.16) plus predicted sORF encoded proteins (Jager et al. (2009), Dar et al. (2016)), and the cRAP list of commonly occurring laboratory contaminants (version 1.0, 2012.01.01). The searches were performed using the Proteome Discoverer software package (version 1.4.0.288) using the SequestHT search algorithm. A tryptic search was performed, (Precursor tolerance 10 ppm, fragment tolerance 0.04 Da, missed cleavages 2). Variable protein modification: oxidation of methionine; fixed modification: cysteine carbamidomethylation. Strict parsimony criteria were applied with a target FDR of 0.01 (1%) applied at peptide level. In addition, proteins required two high confident peptides to be considered as identified.

### Reverse pull-down analysis of immobilized Strep-GlnA_1_ and SUMO-His_6_-sP26

*E. coli* BL21 pRIL containing plasmid pRS375 was grown at 37 °C to a turbidity of 0.6 at 600 nm and then expression of Strep-GlnA_1_ was induced by adding 200 µg/L tetracycline. After 2 h incubation at 37°C, cells were harvest, resuspended in 6 mL W-buffer (100 mM Tris, pH 8.0; 150 mM NaCl; 1 mM EDTA) and disrupted using a French pressure cell at 4,135 × 10^6^ N/m^2^, followed by centrifugation at 20,000 x g for 20 min. The cell free crude extract was incubated with 1 mL W-buffer equilibrated Strep-Tactin^®^ sepharose^®^ matrix (IBA, Göttingen, Germany) incubated for 30 min at 4 °C under slightly swivel. The column was washed with 10 mL W-buffer. Strep-GlnA_1_ was eluted in the presence of 2.5 mM D-Desthiobiotin (IBA, Göttingen, Germany) followed by buffer exchange using an amicon centrifugal filter (30KDa) (Merck; Darmstadt, Germany) according to the manufacturer’s instructions. To prepare cell free crude extract containing SUMO-His_6_-sP26 *E. coli* BL21 pRIL / pRS1229 was grown in 1 L LB at 37 °C to a turbidity at 600 nm of 0.6 and expression of SUMO-His_6_-sP26 was induced by adding 100 µM IPTG, followed by further incubation for 2 h. Cells were harvested and cell free extract was prepared as described above. Approximately 50 mg of crude extract containing SUMO-His_6_-sP26 were mixed with 1 mg purified Strep-GlnA_1_ and incubated with 1 mL Strep-Tactin^®^ sepharose^®^ matrix (IBA, Göttingen, Germany) for 30 min at 4 °C under slightly swivel. Non-binding proteins were subsequently washed from the column with 10 mL W-buffer. Strep-GlnA_1_ and potential interacting proteins were eluted in the presence of 2.5 mM D-Desthiobiotin (IBA, Göttingen, Germany). Aliquots of wash and elution fractions of the reverse co-chromatography were separated by SDS-PAGE according to Laemmli (1970) and the elution fractions were further analyzed by Western Blot analysis as described in Weidenbach et al. (2010) using commercial antibodies directed against the His-tag following the instructions of the manufacture (Qiagen, Hilden, Germany).

### Purification of His_6_-GlnK_1_, His_6_-GlnA_1_ and His_6_-sP26

1 L of the respective *E. coli* BL21 / pRIL strains carrying pRS203 (His_6_-GlnK_1_), pRS196 (His_6_-GlnK_1_) or pRS1245 (His_6_-sP26) were grown in LB-medium at 37°C under rigorous shaking to a turbidity of 0.7 at 600 nm. The cultures were cooled down to 18°C followed by supplementing with 10 µM IPTG and further incubation at 18°C for approx. 18 h. The cells were harvested at 4.000 x *g* and 4 °C for 20 min and resuspended in 20 ml buffer B (50 mM HEPES, 300 mM NaCl, pH 7.5). Cytosolic extracts were generated as described above and the His_6_-tagged proteins purified by metal affinity chromatography and gravity flow using 0.5 ml HisPur™ cobalt superflow agarose (Thermo Fisher Scientific™, Waltham, USA) according to manufacturer’s protocol, eluting proteins at 50 mM imidazole. The imidazole of the elution fraction was removed until the concentration of imidazole was below 1 mM using Amicon^®^ centrifugal filters (0.5 ml size, GlnA_1_: 30 kDa and GlnK_1_: 10 kDa) (Merck KGaA, Darmstadt, Germany). His_6_-sP26 was purified as described above with the following additional steps: The generated cytosolic extract was fractionated by (NH_4_)_2_SO_4_ precipitation prior the metal affinity chromatography. His_6_-sP26 precipitated at approx. 40% to 45% ammonium sulfate. The cytosolic extract was supplemented with 30% (NH_4_)_2_SO_4_ (v/v) and incubated under stirring for 1 h at 8 °C. After centrifugation at 13.865 x *g* and 4 °C for 45 min the respective supernatant was supplemented with additional 15% ammonium sulfate and processed as described. The generated second pellet (precipitated His_6_-sP26) was resuspended in 20 ml Buffer B and dialyzed overnight against buffer B. The protein was purified using HisPur™ cobalt superflow agarose (0.5 ml) (Thermo Fisher Scientific™, Waltham, USA), see above. Finally the His_6_-sP26 protein was separated from contaminating larger proteins using size exclusion 10 kDa Amicon^®^ centrifugal filters (0.5 ml size) and the buffer was exchanged to buffer B without imidazole using 3 kDa Amicon^®^ centrifugal filters (0.5 ml size) (both from Merck KGaA, Darmstadt, Germany).

### Gel electrophoretic separation and Determination of protein concentrations

The purifications of His_6_-GlnA_1_ and His_6_-GlnK_1_ were analyzed using standard 12% SDS-PAGE (Laemmli (1970)), whereas His_6_-sP26 was analyzed using 18% Tricine-SDS-PAGE (Schagger (2006)). For interaction analyzes the gradient 4%-12% Bis-Tris NuPAGE (Thermo Fisher Scientific™, Waltham, USA) according to manufacturer’s protocol were used.

Due to the lack of aromatic amino acids within the small protein His_6_-sP26 the Pierce™ BCA protein assay kit (Thermo Fisher Scientific™, Waltham, USA) was used to determine the concentration of all proteins and was performed according to manufacturer’s protocol. Absorbance was measured at 562 nm by the spectrophotometer Ultrospec 2100 UV/VIS (GE Healthcare™, Chicago, USA). The concentration was calculated by using a BSA standard (50 - 2000 µg/ml).

### Interaction and affinity analysis by Microscale Thermophoresis (MST) using Monolith NT.115

50 µg synthesized sP26 (Davids Biotechnologie, Regensburg, Germany) was fluorescently labeled with the Monolith NT Protein Labeling Kit RED (NanoTemper, Munich, Germany) according to manufacturer’s protocol. 20 nM labeled sP26 in MST buffer (50 mM Tris-HCl pH 7.4, 150 mM NaCl, 10 mM MgCl2, 0.05 % Tween-20) was applied to a dilution series of purified His_6_-GlnA_1_ ranging from nM to µM concentrations and measured using standard treated capillaries (NanoTemper, Munich, Germany), 100% Excitation Power and medium MST-Power.

Purified His_6_-GlnA_1_ was fluorescently labeled using the Monolith Protein Labeling Kit RED-NHS 2nd Generation (NanoTemper, Munich, Germany) with buffer B (50 mM HEPES, 300 mM NaCl, pH 7.5). A degree of labeling (DOL) of 0.95 was achieved and for analysis 20 nM labeled protein, supplemented with 0.05 % Tween20 and 1 mg ml^-1^ BSA, was applied to nM to µM concentrations of purified His_6_-sP26. The measurement was performed using standard treated capillaries, an Excitation power of 20% and high MST-Power.

His_6_-GlnK_1_ was purified and labeled using Monolith Protein Labeling Kit RED-NHS 2nd Generation (NanoTemper, Munich, Germany) (DOL: 0.87). 12 nM labeled His_6_-GlnK_1_ was incubated with purified His_6_-GlnA_1_ (ranging from nM to µM). The interaction was characterized using 50 mM HEPES, 300 mM NaCl, 1 mg ml^-1^ BSA and 0.05% Tween20. The measurements were performed with standard treated capillaries, 20% Excitation power and high MST-Power.

Purified His_6_-sP26 and labeled His_6_-GlnK_1_ were characterized with the same parameters, whereas the dilution series of His_6_-sP26 was ranging from nM to µM concentrations. In general, the proteins were incubated at RT for 5 min prior to loading the capillaries. K_D_ values were calculated using the Nanotemper tool MO.Affinity Analysis. Data of three independent pipetted measurements were analyzed (MO.Affinity Analysis software version 2.3, NanoTemper Technology).

### Determination of glutamine synthetase activity

Glutamine synthetase (GS) activity was determined by using the coupled optical test assay described by Shapiro (Shapiro and Stadtman, 1970), which couples the consumption of ATP by the conversion of ammonium and glutamate to glutamine catalyzed by glutamine synthetase to the oxidation of NADH by lactate dehydrogenase. The test assay was optimized by testing several buffer systems at pH 7 instead of the reported MOPS buffer (TES, HPO_4_^2-^/H_2_PO_4_^-^, imidazole, Tris/HCl and HEPES). The optimized assay buffer used contained 90 mM KCl, 50 mM NH_4_Cl, 30 mM glutamate (pH 7), 50 mM MgCl_2_ and 43 mM HEPES (pH 7) (final concentrations). The test assay was performed as follows: 50 µg purified His_6_-GlnA_1_ were preincubated for 5 min at RT in 0.4 ml freshly prepared concentrated assay buffer, supplemented with 20 µl 14 mM NADH (freshly prepared) (Carl Roth®, Karlsruhe, Germany), 10 µl 100 mM PEP (Carl Roth®, Karlsruhe, Germany), 10 µl of the enzymatic mixture (lactic dehydrogenase (LDH), 900-1400 U, and pyruvate kinase (PK), 600-1000 U) (Sigma-Aldrich, St. Louis, USA) and varying amounts of purified His_6_-sP26 or/and His_6_-GlnK_1_ to a final volume of 0.95 ml. Once the absorbance at 340 nm was constant the reaction was started by adding 50 µl 72 mM ATP (Roche, Basel, Switzerland). The reaction was continuously monitored over a time course of 300 s by a spectrophotometer (Jasco V-550, Pfungstadt, Germany). In case the assays were performed in the presence of 2-oxoglutarate (2-OG), it was supplemented 5 min before the start. The slope was calculated for the linear degression of the absorbance at 340 nm and the Δc was calculated using the Beer-Lambert law.

### Structure analysis using Tycho NT.6

His_6_-GlnA_1_ was analyzed using Tycho NT.6 (NanoTemper, Munich, Germany) by applying a standard capillary (10 µl) with 1.5 mg ml^-1^ enzyme in buffer B. Thermal unfolding profiles were recorded within a temperature gradient between 35 °C and 95 °C. In case the assays were performed with the addition of 2-oxoglutarate (2.5 mM, 5 mM, 7.5 mM, 10 mM and 12.5 mM), it was supplemented respectively 5 min before start.

### Complex analysis using size exclusion chromatography

All proteins were heterologous expressed in *E. coli* and purified as described above: 168.75 µg His_6_-GlnA_1_, 77.5 µg His_6_-GlnK_1_ and 24 µg His_6_-sP26 in different combinations (His_6_-GlnA_1_; His_6_-GlnA_1_ + His_6_-sP26; His_6_-GlnA_1_ + His_6_-GlnK_1_; His_6_-GlnA_1_ + His_6_-GlnK_1_ His_6_-sP26) were incubated together at RT for 5 min in the presence and absence of 5 mM 2-OG prior to loading on the analytical ENrich 650 column (BioRad Laboratories, Inc., Hecules, USA). The column was equilibrated with 50 mM Tris/HCl and 150 mM NaCl and supplemented with 5 mM 2-oxoglutarate when the complexes were preincubated in the presence of 5 mM 2-oxoglutarate and the applied proteins isocratic eluted with the respective buffer with a flow rate of 1 ml min^-1^. 1 ml fractions were collected and concentrated by TCA precipitation (15 % v/v). The pellets were resuspended in 50 mM ammonium acetate for LC-MS/MS analysis. A detailed method description of the LC-MS/MS method is provided in the supplemental Materials and Methods.

## RESULTS

Aiming to gain insight into the physiological role of small protein sP26 in *M. mazei* we addressed the hypothesis that sP26 is directly or indirectly involved in nitrogen regulation. First we examined, whether the archaeal small protein interacts with other *M. mazei* proteins and verified the identified interacting partner glutamine synthetase using various approaches. Finally, the respective consequences due to the interaction were characterized.

### spRNA26 and the corresponding small protein is highly conserved in Methanosarcina strains and upregulated under nitrogen limitation in *M. mazei*

spRNA26 was identified using a dRNA-seq approach and shown to be expressed under N-limitation whereas under N-sufficiency no transcript was detectable (Jager et al. (2009)). It represents an approximately 70 nucleotide (nt) long leaderless RNA encoding a small protein of 23 amino acids, which was verified in the cell extract of cells grown under N starvation by a peptidomics approach, however with lower confidence as the ones reported (Cassidy et al. (2016)). spRNA26 is located within the 316 bp intergenic region between *MM2083* encoding orotidine 5’-monophosphate decarboxylase and the operon encoding the acetyl-CoA-decarbonylase/synthase (ACS) complex. Its promoter (TATA- and BRE-Box) was identified between nt 26 and 40 upstream of the transcriptional start site (+1) (see Fig. 1A). Two further alternative BRE-boxes can be predicted (indicated with a dashed line). The repressor binding motif of the general transcriptional regulator NrpR (Weidenbach et al. (2008)) was not detected. However, the repetitive elements identified upstream of the BRE-box (indicated with grey boxes) might represent a week binding site for the transcriptional activator NrpA which has been identified in the upstream promoter region of the *nif-*operon (Weidenbach et al. (2014)). Multiple alignments of the respective spRNA26 and homologs in other Methanosarcina species showed that the experimentally verified sORF is highly conserved with regard to amino acid as well as the nucleotide sequence (Fig. 1BC). Further, high conservation of the promoter and the 5’up-stream region was identified among the *M. mazei* strains of which four are missing the BRE-box identified for *M. mazei* Gö1 and thus might use one of the alternative BRE boxes (see Fig. 1B). All homologs are predicted to start with valine indicating that a non-canonical start codon is used for translation initiation (GTG). Interestingly, at the third position of the small protein three *Methanosarcina barkeri* strains contain methionine (ATG) in contrast to all other strains encoding valine (GTG) at the third position. This might indicate the presence of an alternative translation start. Unfortunately, the peptide detected by the LC-MS MS analysis starts with the 5^th^ position (lysine) of the predicted sORF (see Fig. 1C, indicated with a box in the sequence logo), thus cannot be used to identify the native start of the small protein. In the *M. thermophila* strains as well as in *M. falvescens* apparently a deletion of nt 33 occurred causing the generation of a translational stop codon subsequently resulting in a short version of sP26 (see Fig. 1C and Fig. S1). Overall, the high conservation of the amino acid as well as the nucleotide sequence argues for a potential physiological role of sP26.

**Fig. 1:**
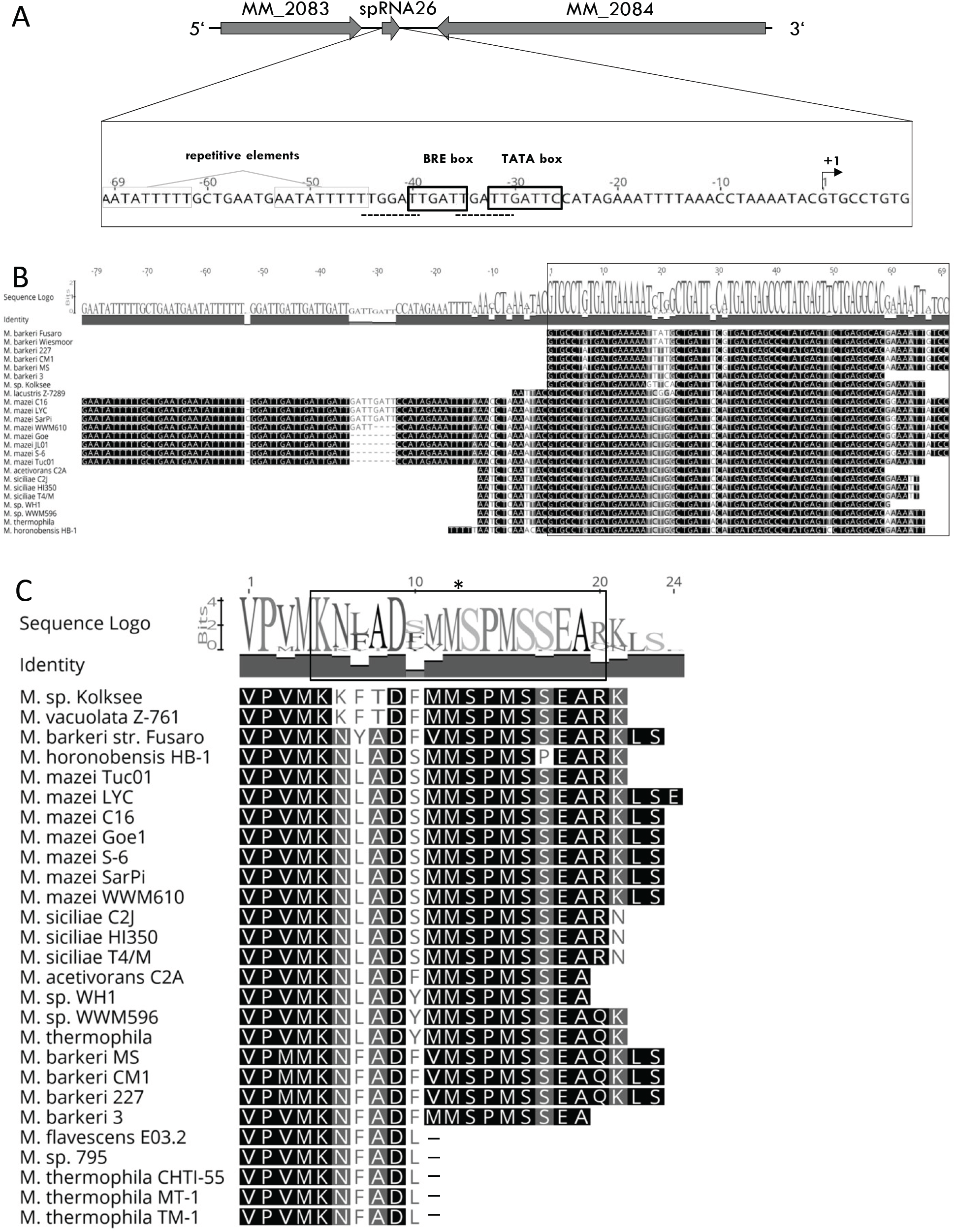
Genomic localization of spRNA26 and homologs in *Methanosarcina* species. (**A**) Location and promoter of spRNA26, predicted BRE and TATA box are indicated as well as alternative BRE boxes with an interrupted line. Additional repetitive elements are boxed (pot. binding sides of NrprA). MM_2083, encoding OMP decarboxylase and MM_2084, encoding γ subunit of acetyl-CoA decarbonylase/synthase. (**B**) Nucleotide alignment (ClustalW) of spRNA26 homologs in *Methanosarcina* species whit the predicted sORF indicated with a box. The similarity is shown in black-grey-white boxes (black symbols 100 % similarity) additional the identity is shown in a grey bar and a nucleotide logo above the nucleotide alignment. (**C**) Amino acid alignment of sP26 homologs in *Methanosarcina* species. The protein sequence was generated by blasting the nucleotide sequence, creating every possible translation frame and aligning every possible sequence to sp26 using ClustalW and further evaluated by hand. The similarity is shown in black-grey-white boxes (black symbols 100 % similarity), the identity is shown in a grey bar and a nucleotide logo above the alignment; *, indicates the peptide identified by LC-MS/MS; -, indicates a stop.

Northern blot and Illumina RNAseq analysis confirmed the reported transcriptional start site (+1) and upregulation of spRNA26 under N limitation (approx. 1.5 to 2.5 fold, see Fig. 2BC). In agreement with the RNAseq data (Fig. 2C), spRNA26 appears to be processed into three shorter fragments (65, 63, and 61 nt) in varying amounts depending on the growth phase and stress conditions (see Fig. 2A).

**Fig. 2:**
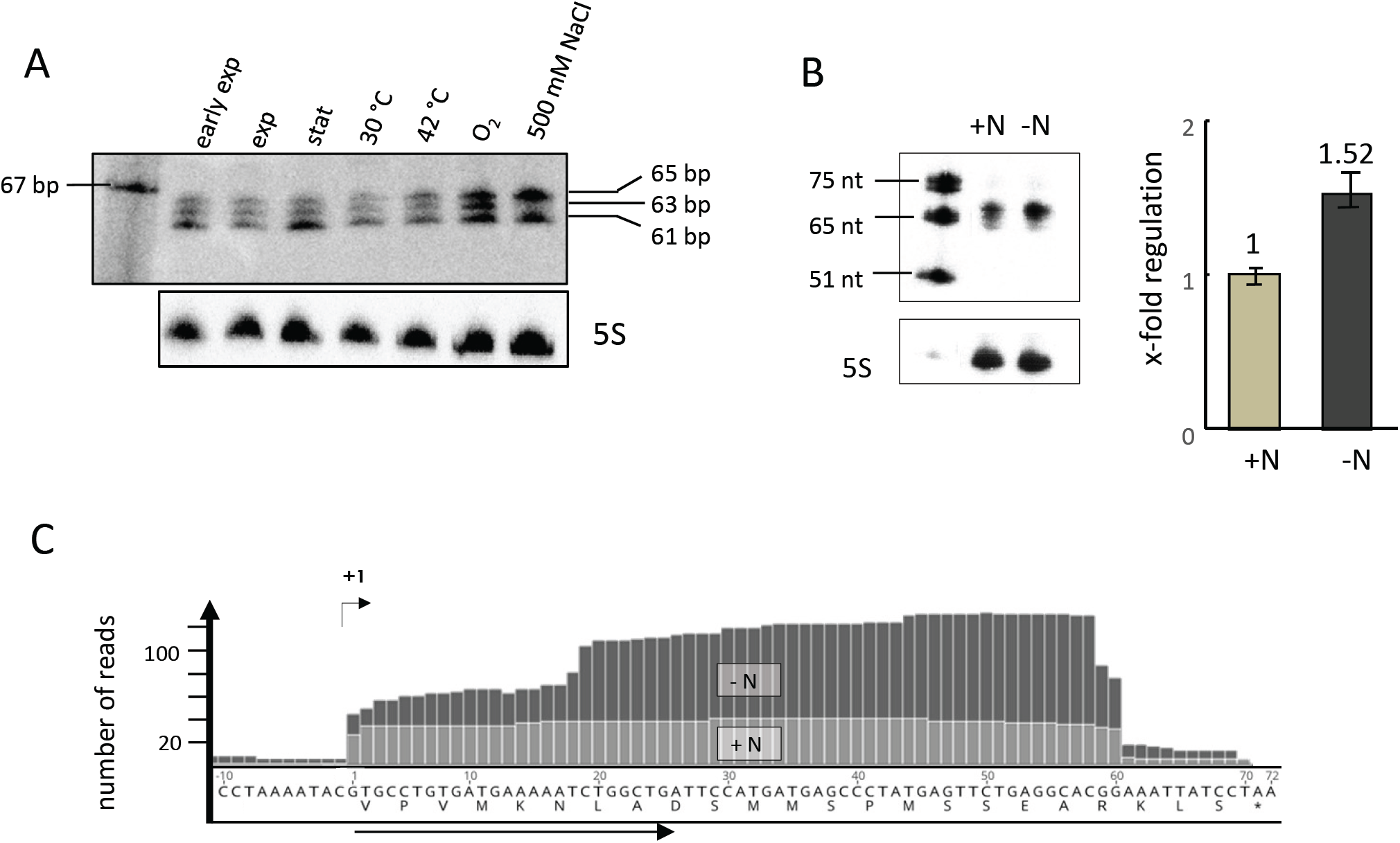
spRNA26 transcription under different growth conditions. (**A, B**) Northern Blot analysis under various conditions. (**A**) Northern Blot analysis was performed as described in Materials and methods using RNA derived from 50 ml cultures grown under nitrogen sufficiency to different growth phases (early exponential, exponential and stationary) and exponential under different stress conditions as indicated. pGEM™ Conventional DNA Digest Marker PR-G1741 (Promega™, Fitchburg, USA) was used as marker. (**B**) *M. mazei* grown exponential under nitrogen sufficiency (+N) and under nitrogen limitation (−N), the regulation was calculated based on eight independent biological replicates, one Northern blot is exemplary shown. (**C**) Differential RNAseq analysis using the Illumina technique was performed with RNA of *M. mazei* grown under nitrogen limitation and sufficiency (exponential growth phase): darker plots: under nitrogen limitation; lighter plots: under nitrogen sufficiency.

To study potential effects of the small protein sP26 *in vivo* the respective sORF was additionally expressed in *M. mazei* from a plasmid under the control of the constitutive promoter *pmcrB* with the ribosome binding site of *mcrB* (pRS1242, plasmid map see Fig. S 2D). Growth analysis under N sufficient and N limiting conditions showed no significant phenotype, growth rates comparable to the control containing the empty pWM321 vector were obtained independent of additional sP26 synthesis (µ (+N)= 0.080 h^-1^ vs. 0.086 h^-1^, µ (−N)= 0.082 h^-1^) (see Fig. S3). Membrane preparations and subsequent Western blot analysis using antibodies raised against synthetic sP26 demonstrated that the majority of chromosomal sP26 is in the cytoplasmic fraction and independent from the nitrogen availability but in a higher oligomeric state or bound to a protein with higher molecular mass (data not shown).

### sP26 is interacting with glutamine synthetase

In order to identify cell proteins directly interacting with sP26 we studied potential complex formation between sP26 N-terminally fused to a SUMO-tag (His_6_-Sumo-His_6_-sP26) and *M. mazei* cell extract proteins by pull-down experiments using affinity chromatography on Ni-NTA agarose for detecting complexes. His_6_-SUMO-His_6_-sP26 was heterologously expressed in *E. coli* (pRS1229, see Fig. S2C) and purified to an apparent homogeneity of 98 % by Ni-NTA affinity chromatography as described in Materials and Methods. Purified His_6_-SUMO-His_6_-sP26 (1 mg) was incubated for 1 h at 4° with approx. 27.5 mg cell extract protein of *M. mazei* cells grown under N limitation. Subsequently, the protein mixture was applied to Ni-NTA agarose (0.5 ml). After washing the chromatography material to remove all cell extract proteins, unspecifically binding to His_6_-SUMO-His_6_-sP26 or the Ni-NTA agarose, His_6_-SUMO-His_6_-sP26 and potential specifically interacting proteins were eluted in the presence of 250 mM imidazole. The respective elution fractions were analyzed by denaturating polyacrylamide gel electrophoresis (SDS-PAGE) und subsequent silver staining. As exemplarily shown in Fig. 3A only one distinct additional protein band corresponding to an approx. 42 kD protein was detected in at least three independent biological replicates. The elution fractions were concentrated and reanalysed on an SDS PAGE stained with Coomassi blue (Fig. 3A, panel 2). The additional protein band was analysed and the presence of glutamine synthetase (GlnA_1_) was demonstrated in two independent biological experiments by LC-MSMS analysis (see Materials and Methods). Unspecific binding of *M. mazei* proteins to the SUMO-fusion protein or the affinity chromatography material was excluded by using purified SUMO-fusion protein and *M. mazei* cell extract (see Fig. 3 A panel 3) and loading *M. mazei* cell extract to the affinity chromatography material followed by elution. To confirm complex formation with GlnA_1_ a revers pull-down was performed. Strep-tagged GlnA_1_ (pRS375) as well as sP26 N-terminally fused to SUMO-tag (pRS1229) were individually heterologously expressed in *E. coli*. Strep-GlnA_1_ was purified to homogeneity by affinity chromatography using Strep-Tactin® sepharose® as described in Materials and Methods. 1 mg purified Strep-GlnA_1_ was incubated with the respective *E. coli* cell extract with His_6_-SUMO-His_6_-sP26 expressed (approx. 50 mg) for 30 min at 4°C followed by affinity chromatography using Strep-Tactin® sepharose®. Strep-GlnA_1_ eluted from the column mainly in elution fraction 2, subsequent Western blot analysis using antibodies directed against the His_6_-tag confirmed the co-elution of His_6_-SUMO-His_6_-sP26 in the Strep-GlnA_1_ elution fractions (see Fig. 3 B).

**Fig. 3:**
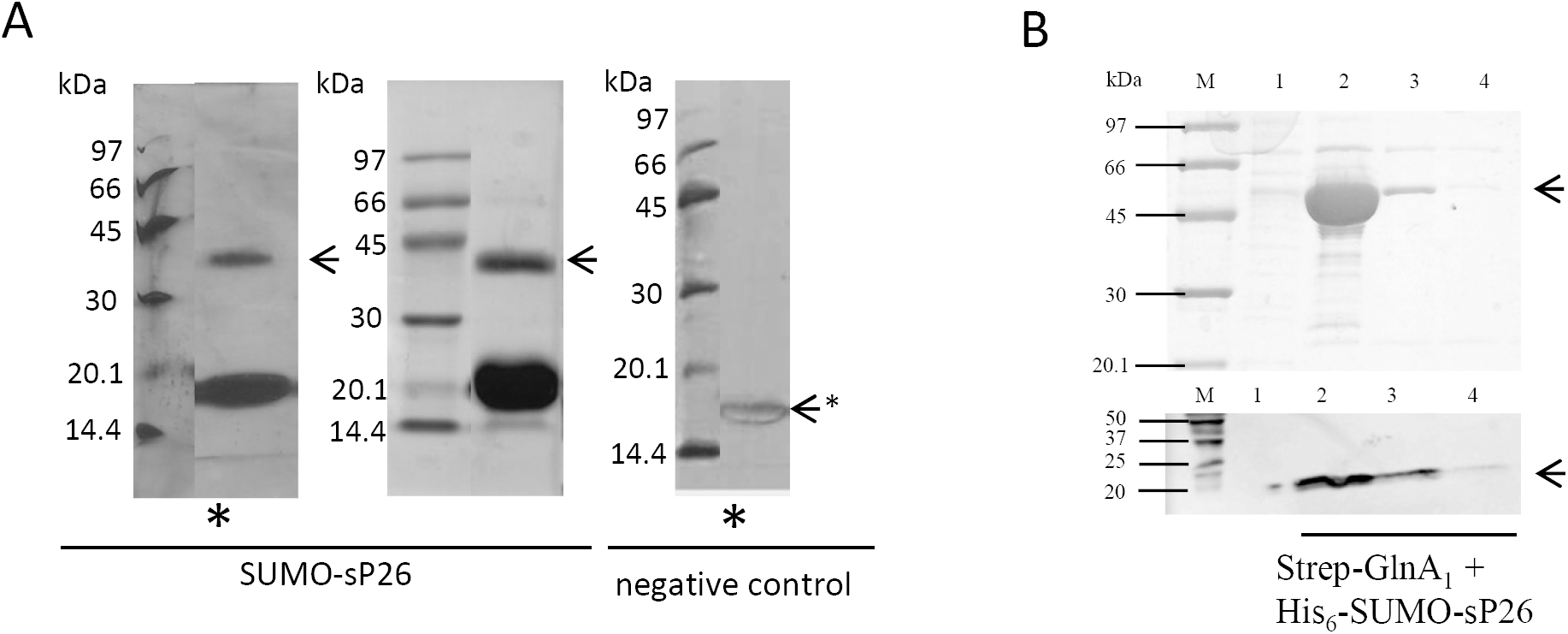
Target identification of sP26 using cell extracts of *M. mazei* grown under nitrogen limitation. **(A) Co-elution analysis of immobilized SUMO-sP26 and *M. mazei* cell extract proteins.** 1 mg purified SUMO-sP26 fusion protein was incubated with 27.5 mg of *M. mazei* cell extract (derived from 1 L culture grown under nitrogen limitation) for 1 h at 4 °C. As control served the SUMO-protein. Proteins were applied to 0.5 ml Ni-NTA agarose. Following the washing steps, SUMO-His_6_-sP26 fusion protein and potentially interacting cell extract proteins were eluted from the affinity material in the presence of 250 mM imidazole and analyzed by SDS-PAGE of the respective co-elution fractions using silver (*) or coomassie staining. The prominent band represents the SUMO-fusion protein (approx. 20 kDa), the potential interacting partner with a calculated size of 42 kDa is indicated with an arrow. (**B) Reverse co-chromatography analysis with purified Strep-GlnA**_**1**_: 1 mg purified Strep-GlnA_1_ protein was pre-incubated with 50 mg *E. coli* cell extract containing overexpressed SUMO-His_6_-sP26 and applied to 1 mL Strep-Tactin^®^ sepharose^®^ matrix (IBA, Göttingen, Germany). After 30 min incubation at 4°C the column was washed followed by elution of Strep-GlnA_1_ and potentially interacting proteins in three elution fractions (1-3), which were analysed SDS-PAGE (upper panel) and Western Blot analysis (lower panel) using antibodies against the His-tag (Qiagen; Hilden, Germany). Lane M (upper panel): Low molecular weight marker (GE healthcare, Freiburg, Germany); Lane M (lower panel): Prescision Plus Protein™ Dual Xtra Standard (BioRad, Feldkirchen, Germany); lane 1, wash fraction; lanes 2-4, elution 1-3.

To obtain further experimental evidence for the interaction between sP26 and GlnA_1_ microscale thermophoresis (MST) analysis was performed and the dissociation constant (K_D_) determined. GlnA_1_ was purified as N-terminal (His)_6_-fusion protein as described by Ehlers et al. (2005b). sP26 was chemically synthesized and nt-647-NHS-labeled as described in Materials and Methods. Interactions between the labeled sP26 (20 nM) and purified His_6_-GlnA_1_ in the range of 0.13 nM to 4.5 µM were analyzed by MST (for details see Materials and Methods). Significant binding was observed in two independent biological experiments, one of which is depicted in Fig. 4A. The K_D_ of 76 +/− 30 nM was calculated based on at least three technical replicates (panel 1), panel 2 shows the respective calculation with one technical replicate. To further verify the interaction, His_6_-sP26 was expressed in *E. coli* (pRS1245, see Fig. S2E), and purified by fractionated ammonium precipitation followed by affinity chromatography on HisPur™ cobalt superflow agarose, and size exclusion using 10 kDa filters as described in detail in Materials and Methods resulting in an appox. 95% purified protein fraction (Fig. 5A). When using purified and nt-647-NHS-labeled GlnA_1_ (20 nM) and purified His_6_-sP26 in the range of 4 nM to 1.4 µM, a K_D_ of 45 µM +/− 14 µM was calculated based on several technical replicates (Fig. 4B, panel 1), which was validated in a second biological replicate, and demonstrates in agreement with the results obtained with the synthesized and labeled sP26 strong interactions between GlnA_1_ and sP26.

**Fig. 4:**
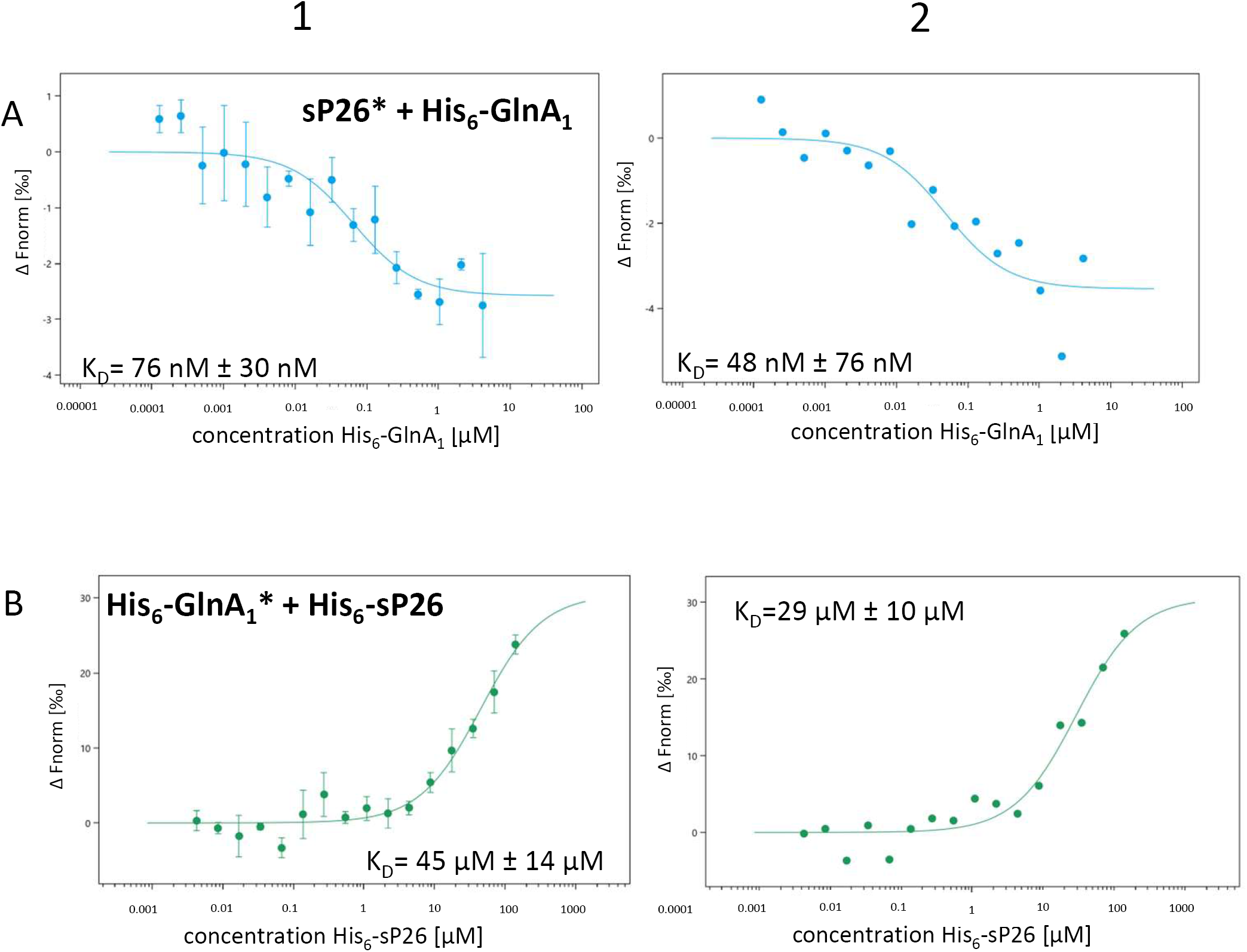
Interaction studies of sP26 and His_6_-GlnA_1_ by Microscale thermophoresis (MST) analysis. (**A**) Synthesized sP26 (Davids Biotechnologie, Regensburg, Germany) (20 nM) was labeled with nt-647-NHS using Monolith Protein Labeling Kit RED-NHS as described in Materials and Methods. The concentration of the purified and non-labeled binding partner His_6_-GlnA_1_ was varied between 0.13 nM to 4.15 µM. After 5 min incubation at RT the samples were loaded to Monolith NT .115 Standard capillaries (NanoTemper Technologies, Munich, Germany) and the MST measurement was performed using Monolith NT. 115 at 100% LED power and medium MST power. A MST-on time of 5 s was used for the analyzed K_D_(His_6_-GlnA_1_): 76 nM ± 30 nM (n=3, error bars represent the standard deviation of one exemplarily chosen biological replicate with three technical replicates (panel 1), one of which is depicted in panel 2). (**B**) Purified His_6_-GlnA_1_ was fluorescently labeled with nt-647-NHS using Monolith Protein Labeling Kit RED-NHS 2nd Generation and kept at a constant concentration of 20 nM. The binding partner His_6_-sP26 was varied between 4 nM – 140 µM and after 5 min incubation at RT the samples were loaded to the standard capillary. MST analysis was performed using Monolith NT .115, standard capillaries, a LED power of 20% and high MST power. MTS-on time of 2.5 s was used for the analyzed K_D_ (His_6_-sP26): 45 µM ± 14 µM (n=3, error bars represent the standard deviation of one biological replicate with three technical replicates, one of which is depicted in panel 2).

**Fig. 5:**
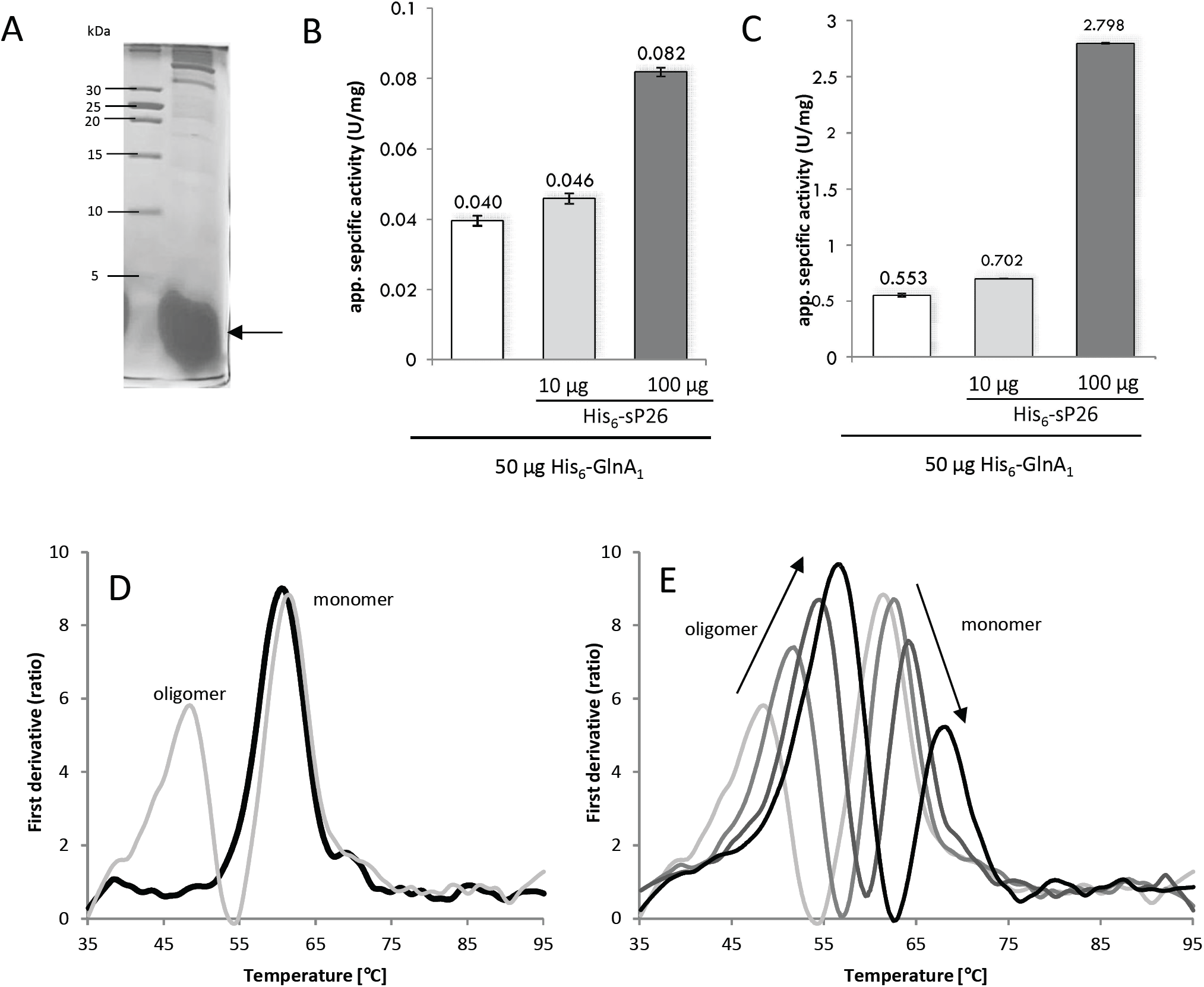
Modulation of His_6_-GlnA_1_ activity by sP26-His_6_ and 2-oxoglutrate (2-OG) and oligomerization state of GlnA_1_. **(A)** Purified His_6_-sP26 was analyzed by SDS PAGE (18 % Tricine gel stained with colloidal comassie as described in Materials and Methods). **(B, C)** Glutamine synthetase activity of purified heterologously expressed His_6_-GlnA_1_ was determined at RT using the coupled optical enzyme assay described by Shapiro and Stadtman (Shapiro and Stadtman (1970)) but in a 50 mM HEPES system in the presence of varying amounts of His_6_-sP26. (**B**) 50 µg purified His_6_-GlnA_1_ and varying amounts of His_6_-sP26 (0, 10 and 100 µg) were preincubated in the absence of 2-OG for 5 min at RT prior to activity analysis. (**C**) The activity was determined under the same conditions but in the presence of 5 mM 2-OG. Depicted are exemplarily data sets from one representative purification batch of GlnA_1_ and sP26. The interaction was confirmed in numerous biological replicates summarized in Table S1. **(D, E)** Structural integrity analysis of purified His_6_-GlnA_1_ dependent on 2-OG (Tycho, NanoTemper, Munich). **(D)** 0.3 mg/ml His_6_-GlnA_1_ were analyzed in the absence (black line) and presence of 5 mM 2-OG (grey line). In the presence of 2-OG two unfolding events occurred the unfolding of the oligomeric form (48°C) and unfolding of the monomer (63°C). **(E)** 0.3 mg/ml His_6_-GlnA_1_ from the same purification were measured with increasing concentrations of 2-OG (5 mM, 7 mM, 10 mM and 12.5 mM - from light grey to dark grey line). With increasing concentration of 2-OG the unfolding events of the oligomeric and monomeric forms shift to higher temperatures indicating a more stable state.

### sP26 effects glutamine synthetase (GlnA_1_) activity in addition to activation by 2-oxoglutarate

Aiming to identify the effects of sP26 on its target GlnA_1_ we evaluated the glutamine synthetase activity in the absence and presence of the small protein. His_6_-GlnA_1_ as well as His_6_-sP26 was purified by affinity chromatography after heterologous expression in *E. coli*. GlnA_1_ activity was determined by using the coupled optical test assay described by Shapiro (Shapiro and Stadtman, 1970), which couples the consumption of ATP by the conversion of ammonium and glutamate to glutamine catalyzed by glutamine synthetase to the oxidation of NADH by lactate dehydrogenase. In course of optimizing the test assay we observed that the buffer system is crucial and significantly effects the specific activity of glutamine synthetase. Using the originally reported MOPS buffer, which was used in the previous report on GlnA_1_ activity (Ehlers et al. (2005b)), significantly lower specific activities were observed (0.026 U/mg) in comparison to other buffer systems (up to 0.082 U/mg, see Fig. S4). HEPES buffer was proven to show the highest specific activities of GlnA_1_ in the absence but also in the presence of the metabolite 2-oxoglutarate (2-OG), which has been demonstrated to significantly enhance GlnA_1_ activity; and represents the internal signal for N starvation in *M. mazei* (Ehlers et al. (2005b). Due to the higher specific activity we consequently exclusively used the HEPES buffer based assay in the following experiments and purified GlnA_1_ in HEPES buffer. In the presence of increasing amounts of purified His_6_-sP26 (10 – 100 µg), which were pre-incubated with 50 µg His_6_-GlnA_1_ (0.95 µM) for 5 min at RT in the test assay before starting by supplementing ATP, glutamine synthetase activity was significantly induced. The positive effect of His_6_-sP26 on GlnA_1_ activity was confirmed by at least five further independent biological replicates using independent protein purifications. In general stimulation of glutamine synthetase activity up to 9-fold were obtained by His_6_-sP26 depending on the respective protein purification (its quality condition) and consequential saturation level (see summarizing supplementary Table S1). One representative set resulting in 2.1-fold stimulation is exemplarily shown in Fig. 5B. Elucidating the effects of His_6_-sP26 on glutamine synthetase activity in the presence of the known activator 2-OG (5 mM), clearly demonstrated that the obtained activating effect of sP26 is independent of the activation by 2-OG and stimulates the activity in addition to the activation due to 2-OG (see Fig. 5C, showing one exemplarily chosen biological activity set). Several independent biological replicates confirmed the additional stimulation of GlnA_1_ activity by His_6_-sP26 in the presence of 2-OG up to 5-fold (see Table S1). For one fraction of purified His_6_-sP26 we additionally tested the positive effects in the presence of 5 mM 2-OG until achieving apparent saturation by His_6_-sP26 (see Fig. S5), indicating that approximately 10 µM His_6_-sP26 is saturating when using 0.95 µM GlnA_1_ in the test assay (calculations based on GlnA_1_ monomers). Consequently, the absolute molecular ratio between His_6_-sP26 and GlnA_1_ is approximately 10:1.

To address the predicted higher oligomeric state of GlnA_1_ in the presence of 2-OG (Ehlers et al. (2005b)) we analysed the conformation and oligomeric state of purified GlnA_1_ by structural integrity analysis using the Tycho system (NanoTemper) as described in Materials and Methods. The analysis clearly showed that in the absence of the metabolite the majority of the purified protein fraction is in the monomeric state (Fig. 5D, black line), whereas the addition of increasing concentrations of 2-OG resulted in increasing amounts of the oligomeric conformation of GlnA_1_ and complementary decreasing amounts of the monomeric fraction (Fig. 5D: 5 mM 2-OG light grey line; Fig. 7E: 5 mM, 7 mM, 10 mM and 15 mM 2-OG from light grey to dark grey). Besides, binding of 2-OG generally resulted in higher stability of both conformations indicated by the significant shift to higher denaturation temperatures (e.g. shifting from 60°C to 70°C for the monomer). Consequently, small variations of the nitrogen status of the respective *E. coli* cultures during independent protein expression might result in varying concentrations of 2-OG bound to purified *M. mazei* GlnA_1_ preparations and thus varying amounts of the oligomeric fraction. Evaluating different independent purifications this variation in the oligomeric fraction might explain the range of stimulation observed by sP26 (Table S1), in addition to the quality of the sP26 purification fractions.

**Fig. 6:**
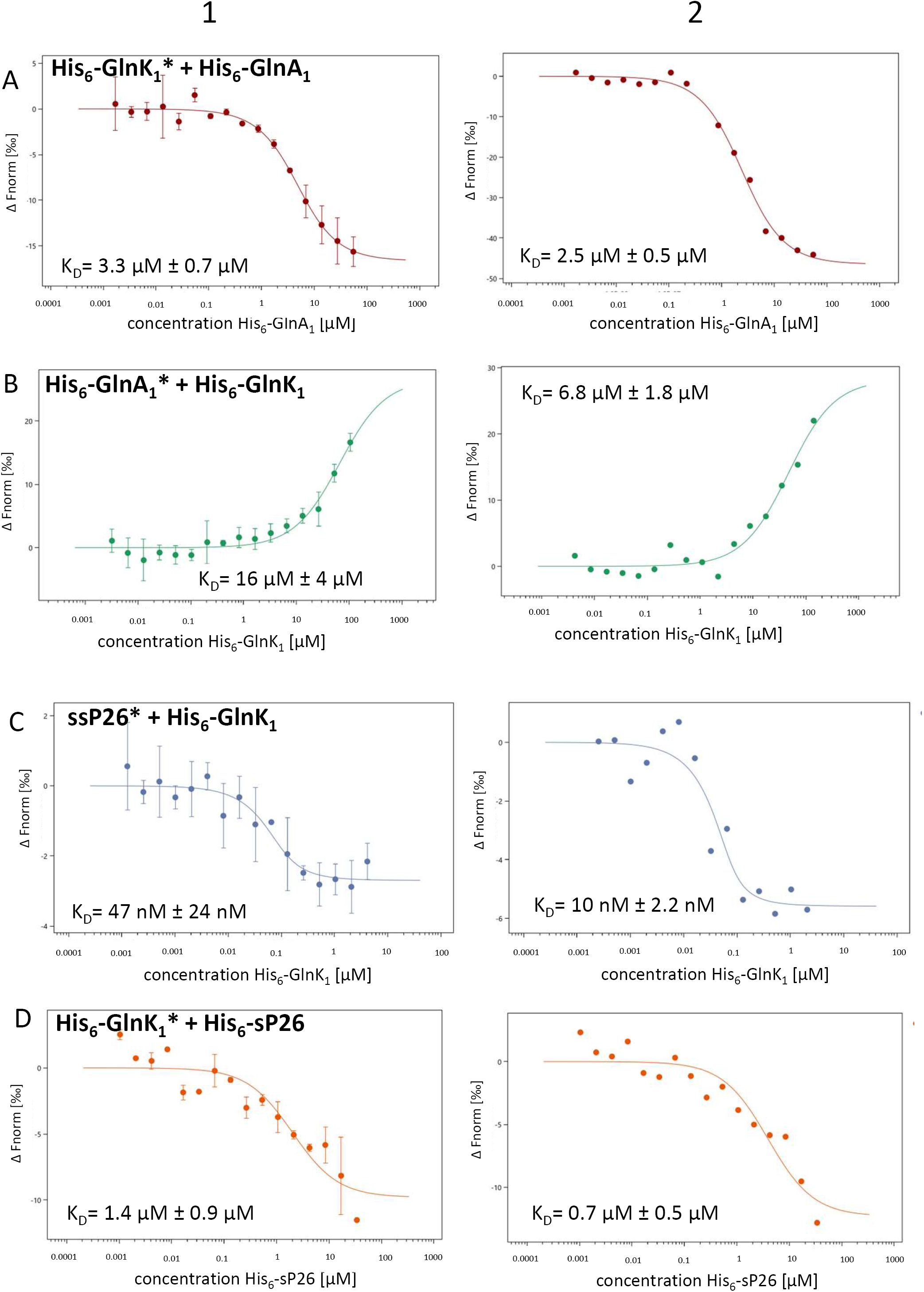
Interaction studies between His_6_-GlnK_1_, His_6_-GlnA_1_ and sP26 by MST analysis: The protein indicated (*) was fluorescently labeled with nt-647-NHS using Monolith Protein Labeling Kit RED-NHS 2nd Generation. The interacting proteins were incubated 5 min at RT prior to loading to standard capillaries (NanoTemper, Munich, Germany). The measurements were performed using Monolith NT.115 with a LED power of 20% and high MST power with a MST-on time of 5 s. **(A)** Purified and labeled His_6_-GlnK_1_^*^ was kept at a constant concentration of 20 nM; purified His_6_-GlnA_1_ was varied between 1.7 nM to 55.5 µM. Calculated K_D_ (His_6_-GlnA_1_): 3.3 µM ± 0.7 µM (with n=2, error bars represent the standard deviation of 1 biological experiment with 2 technical replicates). **(B)** Purified labeled His_6_-GlnA_1_ ^*^ (20 nM) was analyzed with purified His_6_-GlnK_1_ with a concentration range from 6 nM to 105 µM. Calculated K_D_ (His_6_-GlnK_1_): 16 µM ± 4 µM (n=3, error bars represent the standard deviation of 1 biological experiment with 3 technical replicates) **(C)** synthesized sP26 (Davids Biotechnologie, Regensburg, Germany) (20 nM) was labeled with nt-647-NHS using Monolith Protein Labeling Kit RED-NHS. The concentration of the purified binding partner His_6_-GlnK_1_ was varied between 0.13 nM - 4.15 µM. The MST measurement was performed using a Monolith NT. 115 at 100% LED power and medium MST power. Estimated K_D_ (His_6_-GlnK_1_): 47 nM ± 24 nM. (n=2, error bars represent the standard deviation). **(D)** Purified and labeled His_6_-GlnK_1_ ^*^ (20 nM) was analyzed with purified His_6_-sP26 by varying the concentration between 4.3 nM to 34 µM and resulted in an estimated K_D_ (His_6_-sP26): 1.4 µM ± 0.9 µM (n=2, error bars represent the standard deviation). In general, (A-D), out of several biological replicates, one respective biological replicate is exemplarily shown in panel 1 - based on two or three technical replicates, one of which is shown in panel 2.

**Fig. 7:**
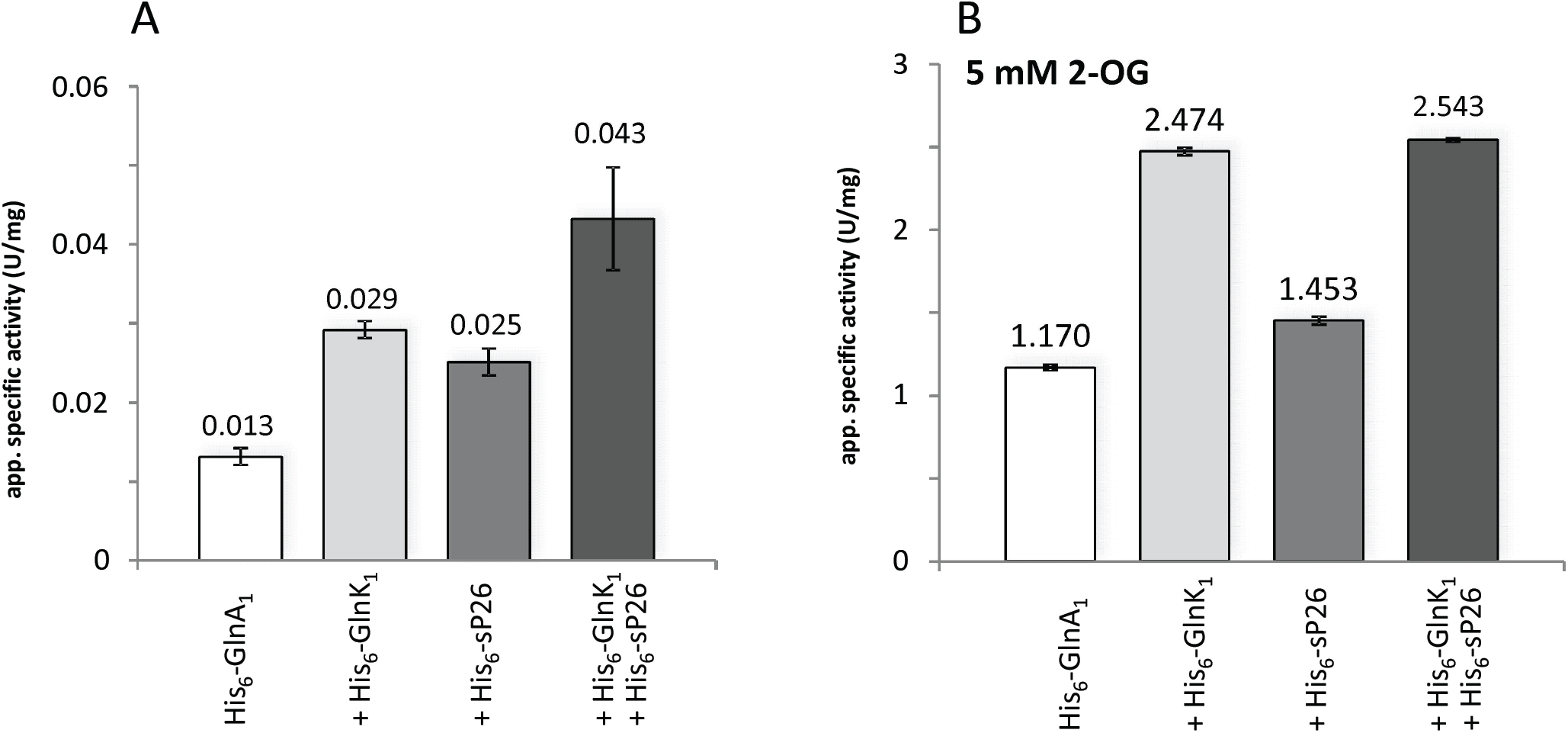
Modulation of His_6_-GlnA_1_ activity by His_6_-sP26 and His_6_-GlnK_1_: Glutamine synthetase activity. Glutamine synthetase activity of purified heterologously expressed His_6_-GlnA_1_ was determined at RT as described in Fig. 7 in the presence of His_6_-sP26 and His_6_-GlnK_1_. (A) 50 µg His_6_-GlnA_1_ was incubated at RT for 5 min, without addition, purified His_6_-sP26 (16 µg), purified His_6_-GlnK_1_ (9.75 µg), purified His_6_-sP26 (16 µg) and His_6_-GlnK_1_ (9.75 µg). (B) The same assays were performed as in (A), but in the presence of 5 mM 2-oxoglutarate. Depicted are exemplarily data from one representative purification batch of His_6_-GlnA_1_, His_6_-sP26 and His_6_-GlnK_1_ (standard deviation of 3 technical replicates). The interaction was confirmed in a second biological replicate (see Table S1).

### sP26 is interacting with GlnK_1_ and stimulates GlnA_1_ activity in addition to activation by GlnK_1_

Previous studies have shown that the PII-like protein GlnK_1_ is modulating the GlnA_1_ activity in *M. mazei* and GlnA_1_ inhibition was predicted due to upshifts in N availability (see Ehlers et al. (2005b)). Consequently, we included GlnK_1_ in our studies evaluating sP26 effects on GlnA_1_ activity. Interaction studies between purified proteins by MST analysis clearly showed and verified the predicted specific interaction between GlnA_1_ and GlnK_1_. Using His_6_-GlnA_1_ and nt-647-NHS-labeled His_6_-GlnK_1_ a K_D_ of 3.3 µM +/− 0.7 µM was calculated (see Fig. 6A) and verified in additional independent biological experiments including a revers labelling experiment (nt-647-NHS-labeled His_6_-GlnA_1_ and His_6_-GlnK_1_, KD = 16 µM +/− 4 µM, see Fig. 6B). For all those experiments, the presence of 5 mM 2-OG did not change the binding affinity (KD-values) (data not shown).

Elucidating potential interactions between GlnK_1_ and sP26 showed in three independent biological experiments that nt-647-NHS-labeled His_6_-GlnK_1_ besides interacting with GlnA_1_ similarly interacts with His_6_-sP26 with high affinities and as well independent of 2-OG (see Fig. 6D, K_D_ = 1.4 µM +/− 0.9 µM). The interaction was further confirmed by an independent reverse labeling experiment analyzing the interaction between His_6_-GlnK_1_ and nt-647-NHS-labeled synthesized sP26 (see Fig. 6C, KD = 47 nM +/− 24 nM). Overall these findings strongly argue that GlnK_1_ modulation has to be taken into further account when analyzing the sP26 effects on GlnA_1_ activity.

Consequently, we included GlnK_1_ in evaluating the enhancing effects of sP26 on GlnA_1_ activity. Aiming to first validate the reported inhibitory effects of GlnK_1_ on GlnA_1_ activity in the new buffer system, we reanalyzed the GlnK_1_ effect in the modified activity assay (see above). Using the originally MOPS based system confirmed the inhibitory effects of GlnK_1_ on GlnA_1_ activity, however changing the buffer system (to NaH_2_PO_4_ or HEPES) resulted in positive effects of GlnK_1_ on GlnA_1_ activity independent of the presence of 2-OG (see Fig. 7 and Fig. S6). Using the HEPES based test assay, we obtained strong evidence in two biological independent replicates that the stimulating effect of sP26 on glutamine synthetase activity is not only independent of 2-OG but also independent and in addition to the stimulating effect of GlnK_1_ (see Fig. 7A, depicting one of two biological replicates). In the presence of 2-OG the additive effects of GlnK_1_ and sP26 were not anymore detectable most likely due to limiting concentrations of NADH and ATP in the test assay (saturation of the activity) (Fig. 7B).

In order to evaluate potential complexes formed between the three proteins, purified His_6_-GlnK_1_ (77.7 µg), His_6_-GlnA_1_ (168.5 µg) and His_6_-sP26 (24 µg) were pre-incubated at RT for 5 min and analysed by size exclusion chromatography (SEC) in the absence and presence of 5 mM 2-OG in the buffer as described in Materials and Methods (ENrich 650 column, BioRad). Analysis of the respective SEC fractions was performed via LC-MS/MS as described in supplemental Materials and Methods.

The results summarized in supplemental figure 7 clearly show that the addition of 2-OG facilitated the identification of both sP26 and GlnK_1_ in the highest molecular weight fraction (>600kDa) besides GlnA_1_, providing further evidence towards the close association of these three proteins.

## DISCUSSION

The recently recognized existence of previously overlooked hidden small proteins in bacterial and archaeal genomes came into focus and of emerging interest. Here, we report on the physiological role of the first archaeal small protein in *M. mazei*. Concordantly we obtained strong evidence that sP26 interacts specifically and with high affinity with the glutamine synthetase (GlnA_1_) in *M. mazei* the key enzyme of ammonium assimilation under N limiting growth conditions. Interaction with sP26 directly effects and stimulates the activity of GlnA_1_ independently and in addition to the inducing metabolite 2-OG and the modulating PII-like nitrogen sensory protein GlnK_1_.

### sP26 stimulates GlnA_1_ activity by direct protein-protein interactions

Ammonium assimilation under N starvation via the glutamine synthetase / glutamate synthase (GS/GOGAT) pathway is one of the major intersections in central N metabolism. Consequently, in all organims synthesis and activity of glutamine synthetase (GS) is strictly regulated by N availability (Reitzer (2003), Reitzer and Schneider (2001), Leigh and Dodsworth (2007)). The GS in Prokaryotes and Eukaryotes can be classified in three different families, GSI, GS II and GSIII, all of which are representing large homo-oligomeric complexes with 8, 10 or 12 monomers. The decameric GSII is present in Eukaryotes, whereas GSI comprising two subtypes, GSI-α and GSI-β and GSIII are found in bacteria and archaea mostly forming dodecamers (Brown et al. (1994)). GSI are typically feedback inhibited by the end products glutamine and AMP. In addition and in contrast to GSI-α, GS of subtype GSI-β are further inhibited in response to N sufficiency by a specific covalent modification of a tyrosine residue near the active side (adenylation by an adenlylyltransferase) (Brown et al. (1994), Smith et al. (1997), Cohen-Kupiec et al. (1999)). In *M. mazei* the essential GS, GlnA_1_, does not exhibit an adenylylation site nor have homologues of adenylyltransferase been identified in the *M. mazei* genome (Deppenmeier et al. (2002), Cohen-Kupiec et al. (1999)) excluding post-translational regulation of *M. mazei* GS in response to N by covalent modification (Brown et al. (1994), Smith et al. (1997), Cohen-Kupiec et al. (1999)). Accordingly, GlnA_1_ represents a GS of the GSI-α subdivision. The gene encoding the essential GS in *M. mazei* (*glnA*_*1*_) is under the direct transcriptional control of the global nitrogen repressor NrpR and thus exclusively transcribed under N limitation (Weidenbach et al. (2008), Jager et al. (2009)). In addition, expression of *glnA*_*1*_ is post-transcriptionally regulated by sRNA_154_, which significantly stabilizes *glnA*_*1*_ transcripts under N limitation (Prasse et al. (2017)). In respect to post-translational N dependent regulation of GlnA_1_ activity, we have shown in the past that GlnA_1_ activity is directly stimulated by the cellular metabolite 2-OG (Ehlers et al. (2005b)). Due to sever reduction of glutamate dehydrogenase activity under N limitation the cellular concentrations of 2-OG increase drastically, consequently the cellular 2-OG concentration is considered to reflect the internal signal for N starvation (Ehlers et al. (2005b)). Perceiving the signal for N starvation as a result of increasing intracellular 2-OG concentrations is also proposed for other autotrophically growing microorganisms as reported e.g. for cyanobacteria (Irmler et al. (1997), Forchhammer (2004), Leigh and Dodsworth (2007)), and has been demonstrated for the post-translational modulation of nitrogenase activity by PII like proteins (NifJ_1_ and NifJ_2_) in response to N availability (switch on, switch off mechanism by direct protein interaction) in *Methanococcus maripaludis* (Dodsworth and Leigh (2006) and Dodsworth and Leigh (2007)). As known for other methanoarchaea, the cellular 2-OG concentration in *M. mazei* also mediates the N status to the transcriptional regulatory machinery. Under N limitation and high internal 2-OG concentration, 2-OG binds to the global repressor NrpR, significantly lowering the binding affinity of NrpR to its respective operator. Consequently, NrpR leaves the operator and transcription can be initiated by RNA polymerase (Weidenbach et al. (2008)). This further emphasizes the central role of 2-OG in perceiving and transmitting the internal N status. In addition, a PII-like protein, GlnK_1_, allows fine tuning control of the glutamine synthetase activity under changing N availabilities and was predicted to inhibit GlnA_1_ activity due to an ammonium upshift after a period of N limitation (Ehlers et al. (2005b)).

Realizing that a wealth of small proteins exists in *M. mazei* identified by systematic global screens for translated small proteins (Cassidy et al. (2019), Cassidy et al. (2016)), now unraveled that GlnA_1_ regulation in *M. mazei* is even more complex and argues for a mechanistically novel posttranslational regulation by a small protein. Using different approaches, we obtained conclusive experimental evidence that the small protein sP26, which is upregulated under N limitation, interacts with GlnA_1_ in *M. mazei* and simultaneously increases its enzyme activity independently and in addition to the stimulating metabolite 2-OG and modulating GlnK_1_. (i) Pull-down experiments demonstrated that purified *M. mazei* His_6_-SUMOsP26 directly interacts with GlnA_1_ in cell extracts, which was verified by reverse pull-down, demonstrating co-elution of SUMO-His_6_-sP26 when purifying Strep-GlnA_1_ (Fig. 3). (ii) High binding affinity between sP26 and GlnA_1_ were observed using MST analysis (including reverse labelling, see Fig. 4). This specific binding was not depending or affected by 2-OG. (iii) sP26 significantly activates glutamine synthetase activity reaching saturating effects when the molecular ratio was approximately 10:1 (Fig. 5 BC, Fig. S5). (iv) This activity stimulation was independent and additive to the 2-OG activation and GlnK_1_ modulation (Fig. 7). (v) First SEC analysis provided evidence towards close association of GlnA_1_, GlnK_1_ and sP26 (see Fig. S7).

Interestingly, in cyanobacteria, where 2-OG is also reflecting the internal N status, small proteins modulating GS activity have been reported several years ago (Garcia-Dominguez et al. (1999), Garcia-Dominguez et al. (2000)). However, these proteins are in the range of 7 and 17 kDa and have been demonstrated to inhibit GS activity by protein-protein interaction due to an ammonium up-shift after N starvation (Garcia-Dominguez et al. (1999)) (see Fig. 8C). Those so-called inactivating factors (IF7 and IF17) are expressed under N sufficiency combining transcriptional regulation but also post-transcriptional regulation by regulatory RNAs, an asRNA (NsiR4) in case of IF7 (Klahn et al. (2015)) and a recently discovered glutamine-binding riboswitch in case of IF17 (Klahn et al. (2018)).

**Fig. 8:**
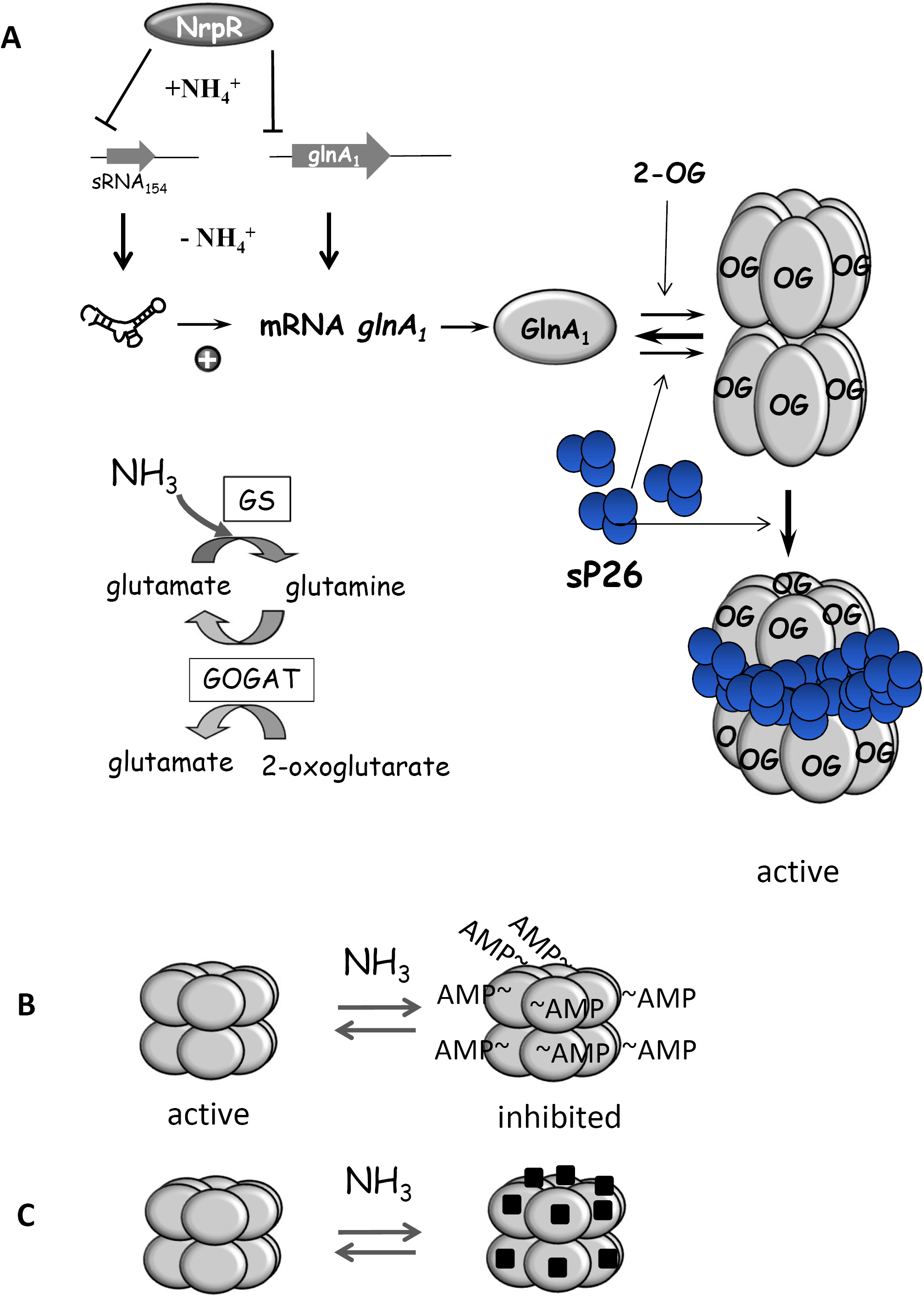
Hypothetical model of GlnA_1_ in *M. mazei*. (**A**) Transcriptional, post-transcriptional and post translational regulation of GlnA_1_. sP26 is proposed to effect GlnA_1_ oligomeric structure in addition to 2-OG by allowing higher stability of dodecamers or tighter dodecamers. sP26 might in addition directly or indirectly effect the GlnA_1_ coupling with GOGAT, resulting in transferring the amino-group of glutamine onto 2-OG, as depicted in the insert. (**B**) Regulation of GS by covalent modification due to N sufficiency (adenylylation) as reported for Proteobacteria. (**C**) Regulation of GS by inhibitory factors in response to N-sufficiency in cyanobacteria.

### sP26 interacts as well with GlnK_1_ in a 2-OG independent manner

As already observed by Ehlers et al. (2005b) the PII protein GlnK_1_ is able to modify GlnA_1_ activity. Direct protein interaction has been exclusively shown by pull-down approaches without quantification. Here, we succeeded to demonstrate by MTS analysis that GlnK_1_ binds to GlnA_1_ with high affinity, and further excluded that the interaction is dependent on 2-OG (Fig. 6). In contrast however, to the previous report (Ehlers et al. (2005b)) we obtained strong evidence that GlnK_1_ is activating GlnA_1_ activity when the enzyme activity is determined in a HEPES buffer based activity assay (Fig. 7). The observation of contradictory effects of GlnK_1_ on GlnA_1_ activity depending on the buffer systems suggests that the buffer system is highly influencing/effecting GlnA_1_ activity. We can only speculate that the different buffer systems, salt and / or differences in the pK_S_ might affect the interaction between GlnA_1_ and GlnK_1_, and or the oligomeric conformation of GlnA_1_. We clearly demonstrated that sP26 also binds to GlnK_1_ with high affinity and in a 2-OG independent manner (Fig. 6CD). Thus, in future it will be important to elucidate the complex formation between all three proteins in the presence and absence of 2-OG in detail and determine the respective ratio as well as the oligomeric conformation of GlnA_1_.

### Hypothetical model for post-translational regulation of GlnA_1_ by sP26

Since strong interactions between sP26 and GlnA_1_ have been observed and it is highly unlikely that sP26 displays a catalytic activity for e.g. covalent modification of GlnA_1_, we propose that the obtained sP26 dependent induction of GlnA_1_ activity is most likely due to effects on GlnA_1_ oligomeric structure or stability. Moreover, sP26 might in addition directly or indirectly effect GlnA_1_ interaction with GOGAT. Potential effects on the GlnA_1_ /GOGAT interaction by sP26 is particularly attractive, since we have detected peptides derived from both large subunits of GOGAT in several pull-down experiments in the LC-MS/MS analysis.

Based on our findings we hypothesize the following model depicted in Fig. 8A. In the presence of 2-OG, GlnA_1_ nearly exclusively expressed under N starvation forms higher oligomeric structures (dodecamers) resulting in GlnA_1_ activation. However, the dodecamers are not very stable. Only in the additional presence und upon interaction with sP26, which is induced under N starvation, GlnA_1_ dodecamers change into a more stable (tighter) complex and consequently induce GlnA_1_ activity in addition to 2-OG. This hypothesized change in the dodecameric structure (two hexameric ring structure face to face) has to be proven in future studies e.g. by NMR analysis and in combination with hydrogen-deuterium exchange mass spectrometry (HDX-MS) to identify the interaction sites at the protein surfaces. Moreover, since sP26 is highly conserved in the Methanosarcinas, it is attractive to speculate that for other methanoarchaeal glutamine synthetases modulations by the sP26 homologs might occur.

## Acknowledgements

We thank the members of our laboratory for useful discussions on the manuscript, as well as Cornelia Goldberg for technical assistance. This work was supported by the German Research Council (DFG) priority program (SPP) 2002 ‘Small proteins in Prokaryotes, an unexplored world’ [Schm1052/20-1, TH 872/10-1, SCHW701/21-1]

## Supplemental Material

**Supp. Fig. 1:**
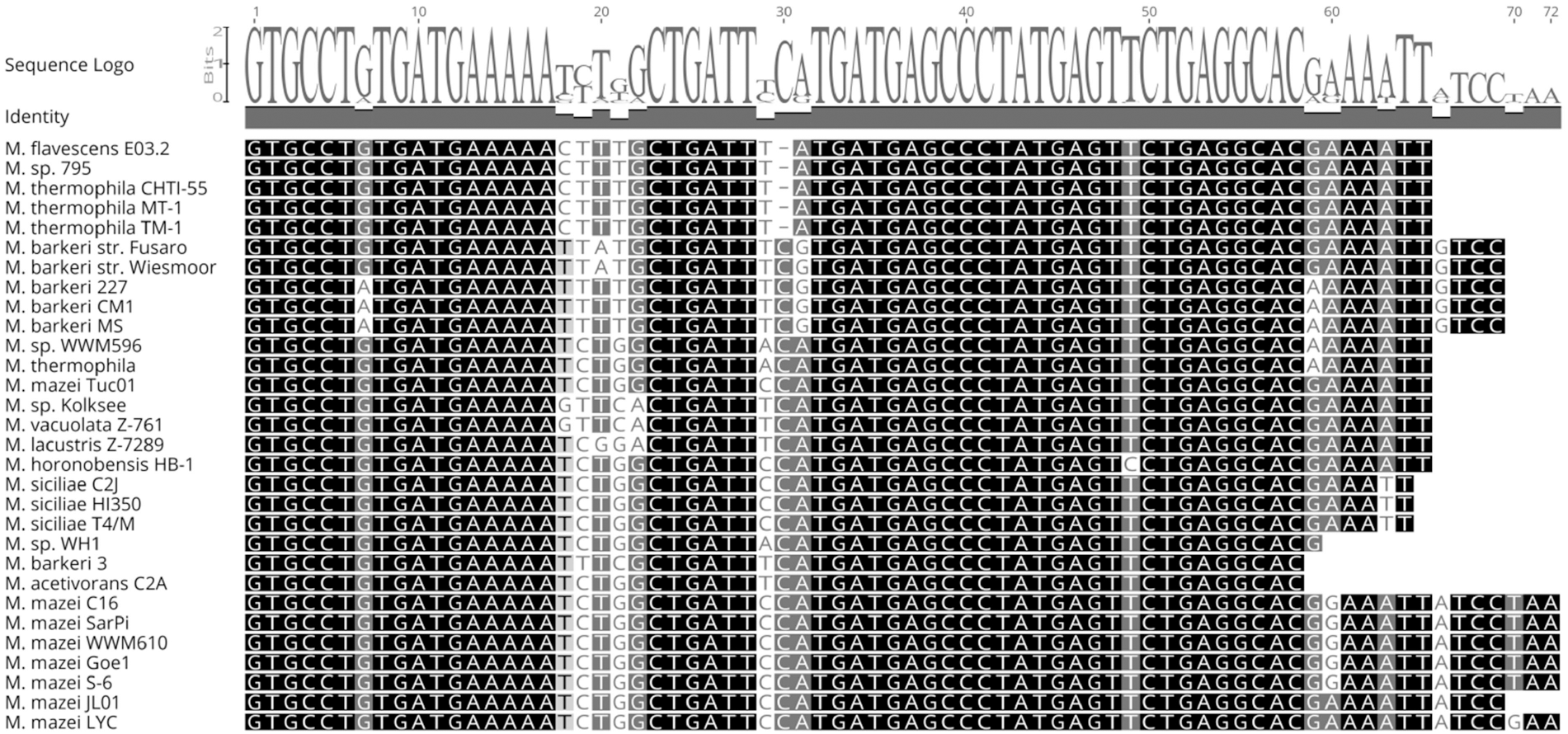
Alignment of homologs of sORF26 in *Methanosarcina* species (ClustalW). The similarity is shown in black-grey-white boxes (black symbols 100% similarity) additional the identity is shown in a grey bar and a nucleotide logo above the nucleotide sequence.

**Supp. Fig. 2:**
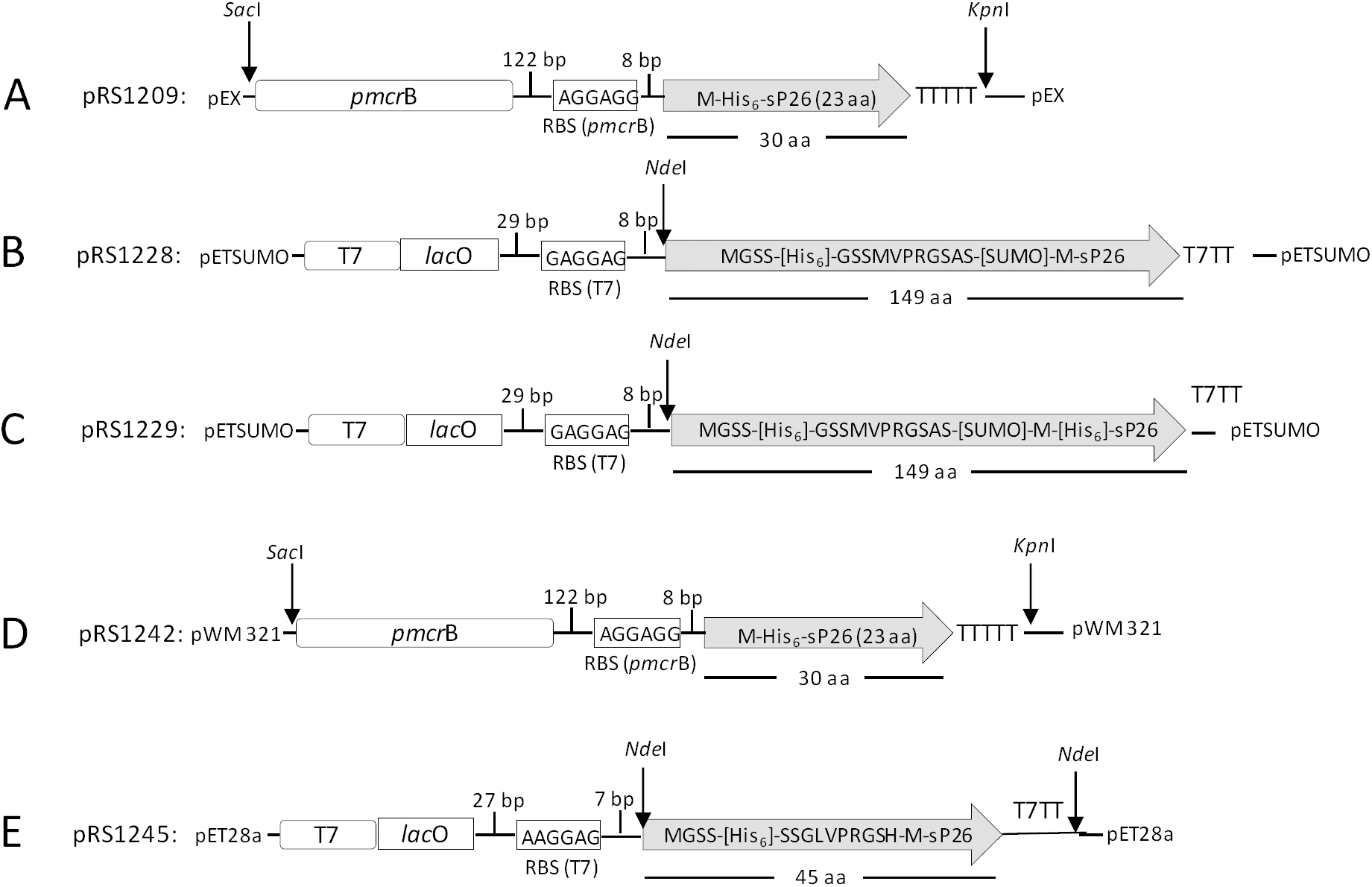
Schematic plasmid maps of expression vectors: (A) pRS1209 containing His_6_-sORF26 in pEX (gene synthesis, Eurofins Scientific, Nantes). (B) pRS1228 containing SUMO-sP26 for heterologous expression in *E. coli.* (C) pRS1229 containing SUMO-His_6_-sP26 for heterologous expression in *E. coli.* (D) pRS1242 containing His_6_-sP26 for expression in *M. mazei.* (D) pRS1245 containing His_6_-sP26 for heterologous expression in *E. coli.* TTTT, transcriptional terminator for *M. mazei;* T7TT, transcriptional terminator T7 RNA polymerase.

**Supp. Fig. 3:**
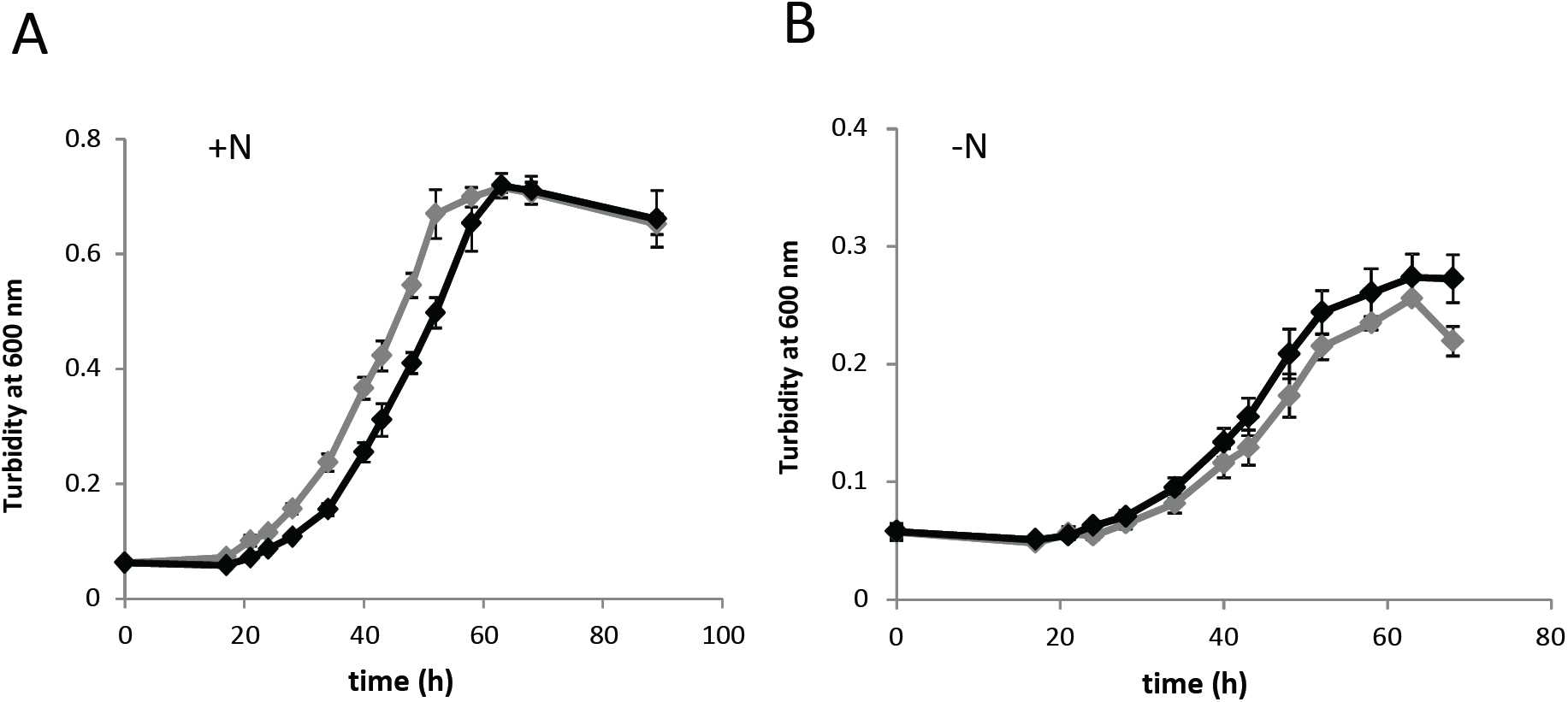
Characterization of *M. mazei* mutant strain overproducing sP26. sP26 was additionally produced in *M. mazei* from pRS1242 containing sORF26 under the control of the *pmcr*B promoter (see Fig.S2). *M. mazei* containing pWM321 served as vector control. Both strains were grown on methanol in the presence of puromycin at 37 °C (50 ml) and growth was monitored. The standard deviation of three independent growth experiments is depicted. (A) Grown in 50 ml under nitrogen sufficiency, (B) under nitrogen limitation with molecular nitrogen as the sole nitrogen source. grey line, *M. mazei* containing pWM321 (vector control); black line, *M. mazei* containing pRS1242 (overproducing sP26); standard deviation is shown and calculated based on nine biological replicates.

**Supp. Fig. 4:**
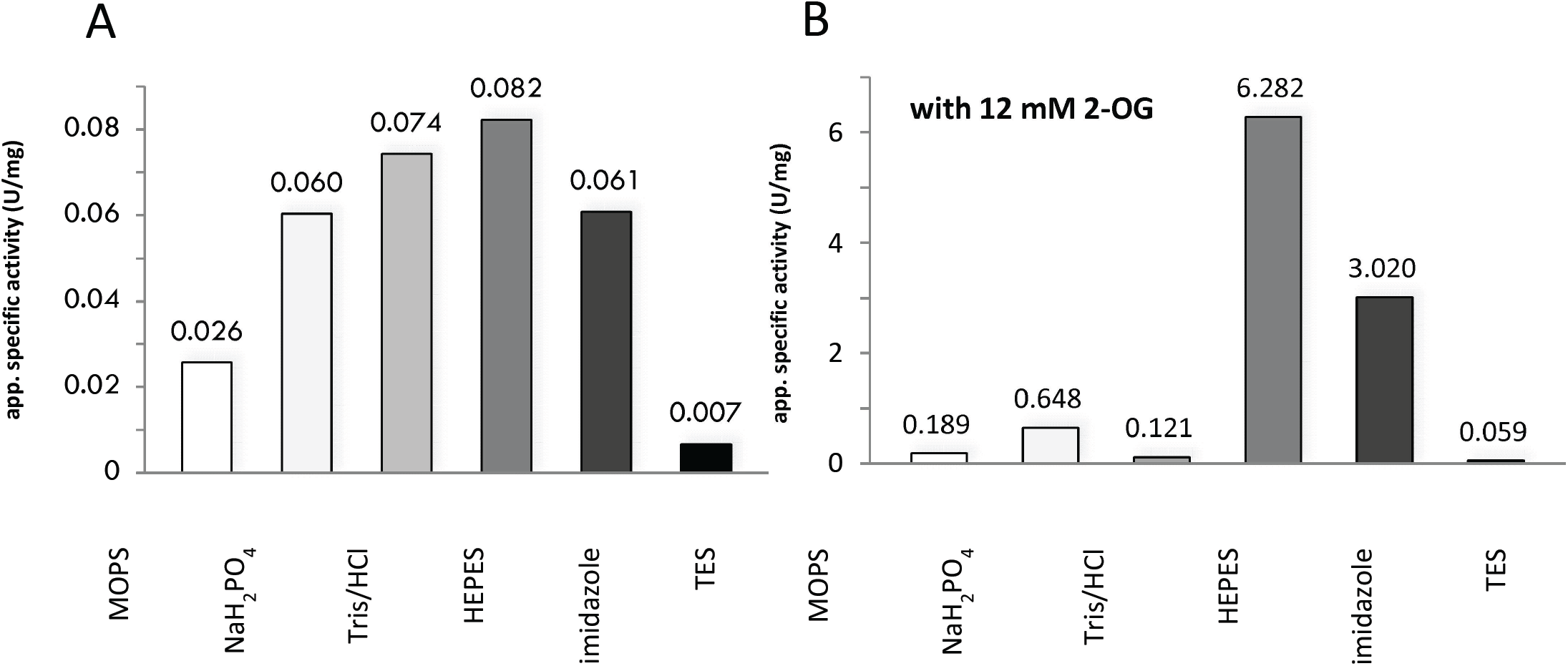
Determination of GlnA_1_ activity in different buffer systems. (A) 50 µg purified His_6_-GlnA_1_ was analyzed by the coupled enzyme assay using different buffer systems. The respective buffers indicated (50 mM, pH 7) contained 300 mM NaCl. (B) the same assays were performed in the presence of 12 mM 2-oxoglutarate.

**Supp. Fig. 5:**
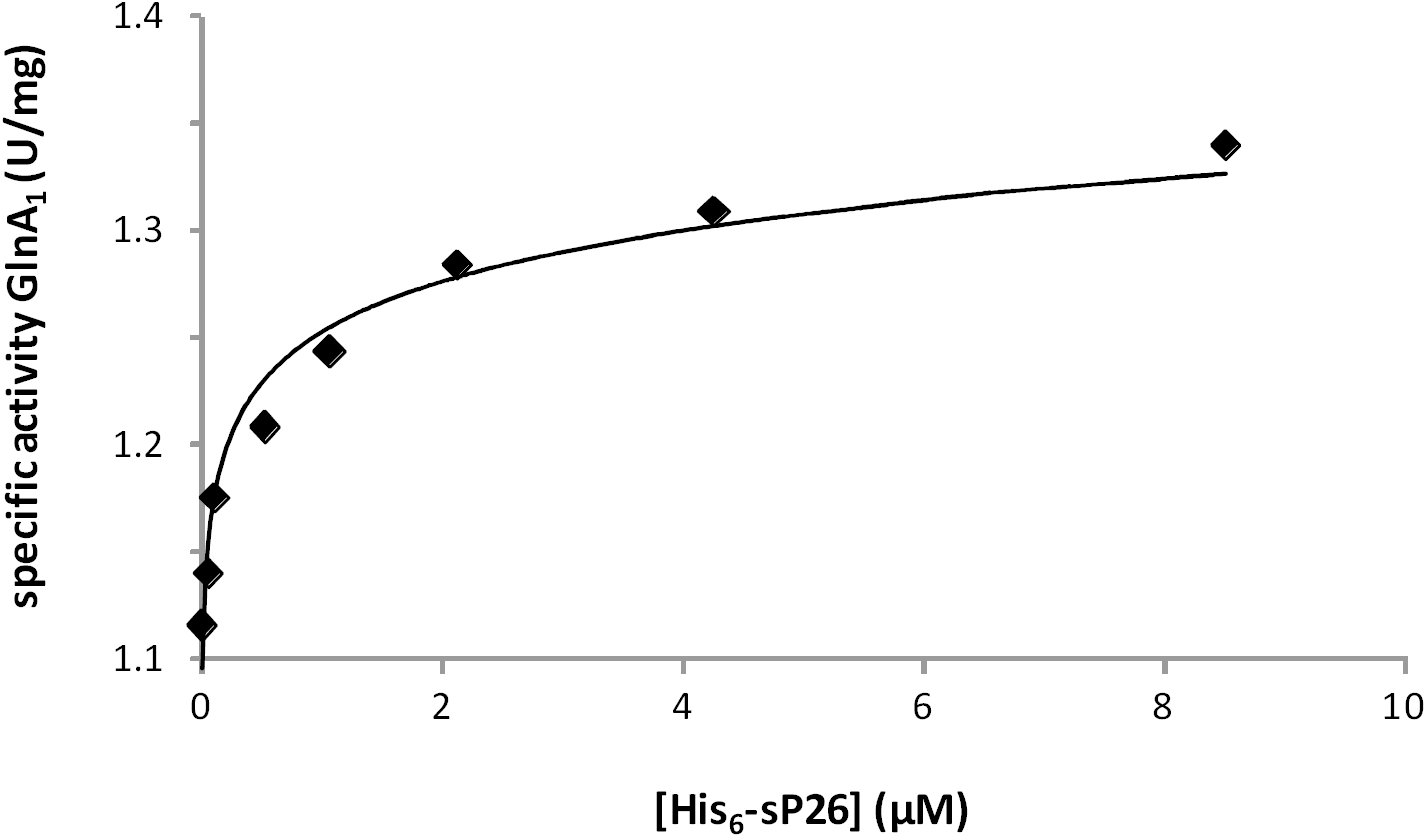
Saturation of the positive effect of sp26 on GlnA_1_ activity. The activity of purified 50 µg His_6_-GlnA_1_ (0.95 µM monomer) in the presence of 5 mM 2-oxoglutarate was measured as described in Fig. 5. Various amounts of purified sP26-His_6_ were added and preincubated for 5 min prior the measurement. All data points were generated using the same His_6_-GlnA_1_ and His_6_-sP26 purification batch.

**Supp. Fig. 6:**
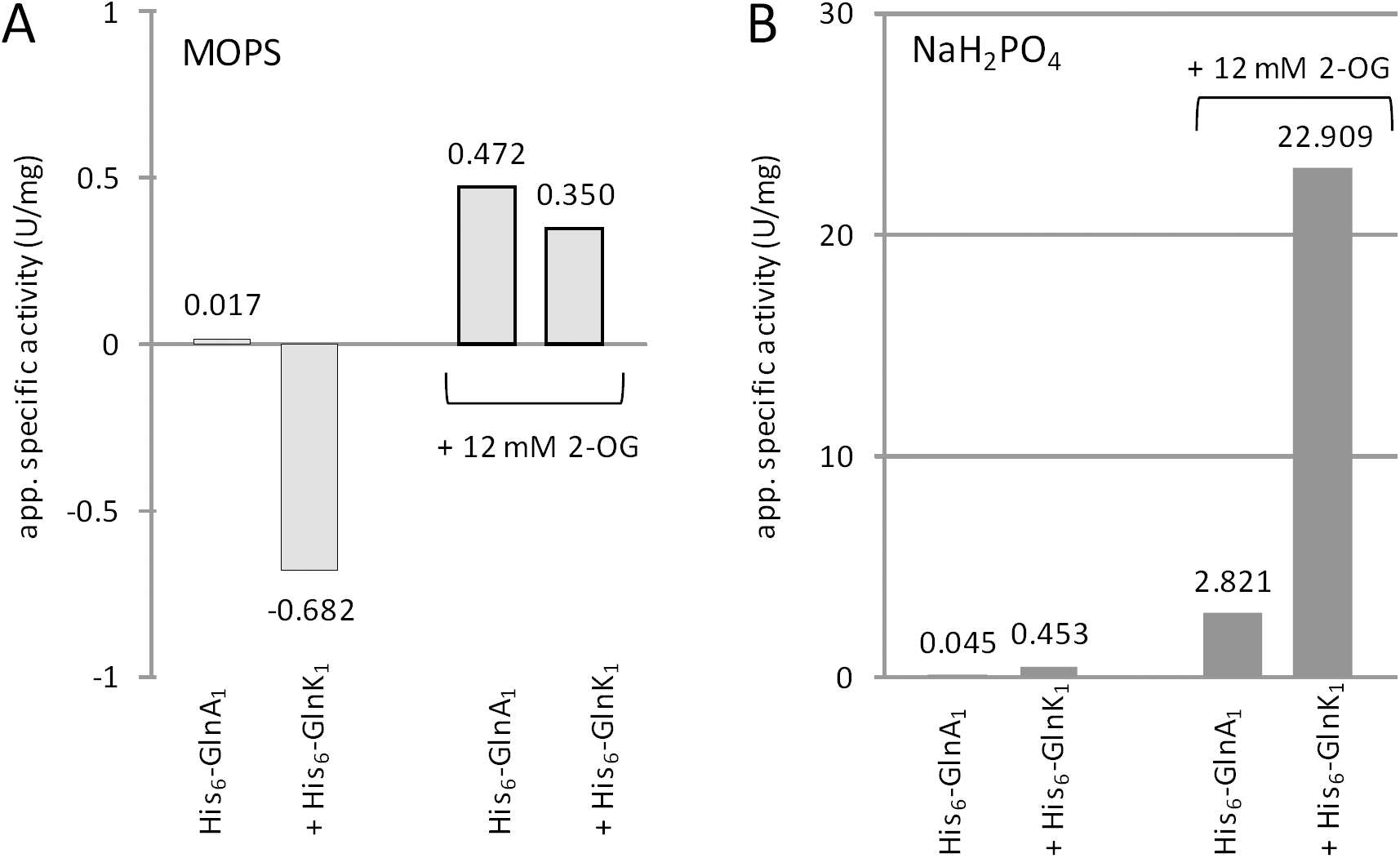
GlnA_1_ activity modulated by GlnK_1_ depending on different buffer systems: 50 µg purified His_6_-GlnA_1_ was analyzed by the coupled enzyme assay with purified His_6_-GlnK_1_ in **(A)** 50 mM MOPS or **(B)** 50 mM sodium hydrogen buffer; the same assays were repeated in the presence of additional 12 mM 2-oxoglutarate.

**Supp. Fig. 7:**
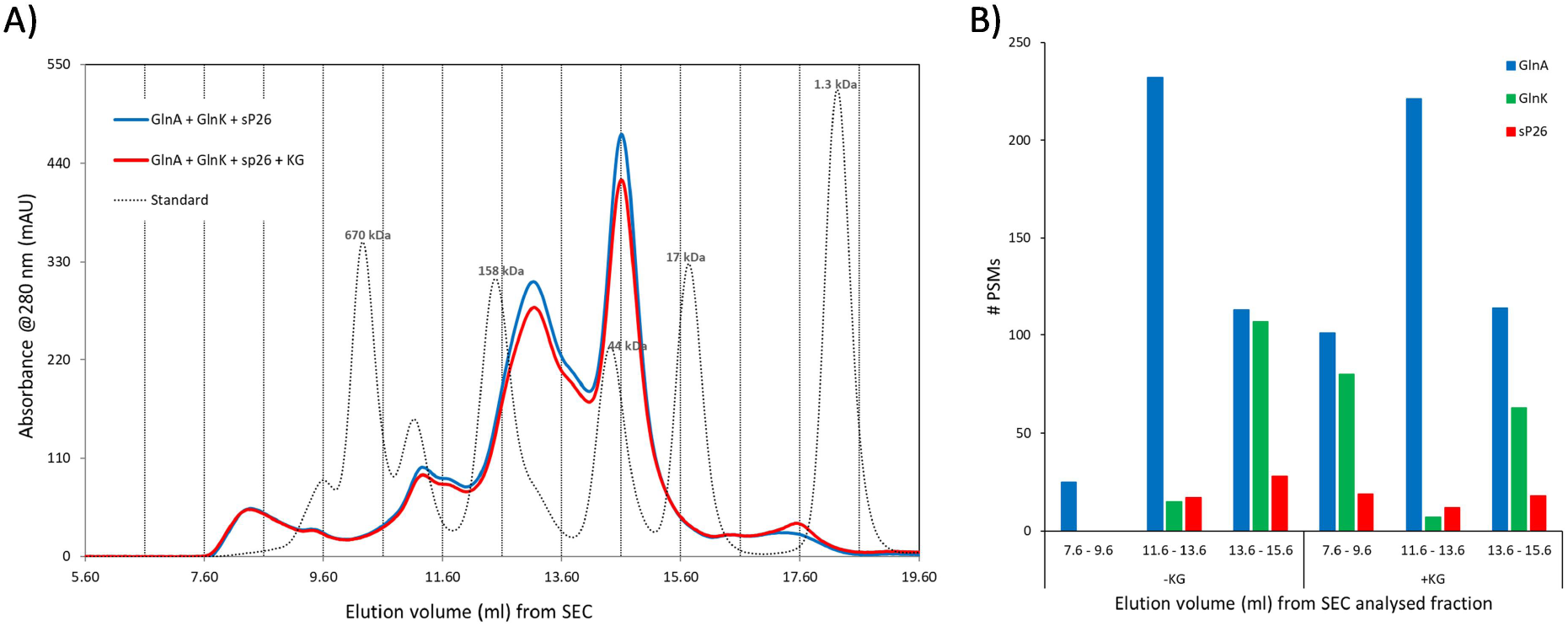
Gel filtration and LC-MS/MS analysis of complex formation between purified *M. mazei* sp26-His6, His6-GlnA1, and His6-GlnK1 proteins in the absence or presence of 2-oxoglutarate. **A)** Gel filtration analysis was performed on a ENrich 650 column (BioRad) using 50 mM Tris/HCl with 150 mM NaCl and a pH 7.5 as buffer system in the presence and absence of 5 mM 2-oxoglutarate (KG) and a flow rate of 1.0 ml/min. Proteins were detected by monitoring the absorbance at 280 nm. Calibration of the column was performed using the gel filtration mass standard (BioRad Laboratories) containing thyroglobulin (670 kDa), IgG (150 kDa), myoglobulin (44 kDa), ovalbumin (17 kDa), and vitamin B12 (1.35 kDa). Elution volumes of standard proteins are indicated. 168.75 µg His_6_-GlnA_1_, 77.5 µg His_6_-GlnK_1_ and 24 µg His_6_-sP26 were pre-incubated in a total volume of 250 µl for 5 minutes at RT before injection. **B)** Protein identification for pooled SEC fractions was performed via LC-MS/MS analysis. The peptide spectral match (PSM) counts for the three proteins of interest (His_6_-GlnA_1_, His_6_-GlnK_1_, and His_6_-sP26) are displayed indicating in which fractions the proteins were identified and providing a estimation of the potential protein abundance. Co-incubation of the proteins with 2-oxogluterate allowed for the identification of both sP26 and GlnK in the highest molecular weight fraction (7.6-9.6 ml).

### Supplemental Materials and Methods for Proteomic analysis for SEC fractions

#### Proteomics analysis of SEC fractions

SEC fractions were precipitated before being suspended in 100 mM ammonium acetate. Aliquots (10 µl) from each fraction were made up to a volume of 60 µl with 100 mM ammonium bicarbonate (ABC) buffer (pH 7.4). The proteins were reduced with dithiothreitol (10 mM, 56°C, 1 hr), and subsequently alkylated in the presence of chloroacetamide (50 mM, room temperature, 30 min). Enzymatic digestion of was performed overnight at 37°C by the addition of sequencing grade trypsin (Promega) (100 ng per sample) in 100 µl of ABC buffer. The samples were acidified via the addition of 10 µl of 10 % TFA to stop the digestion before being dried down under vacuum (Concentrator plus, Eppendorf). The dried peptides were stored at −20°C prior to LC-MS analysis. On the day of LC-MS analysis the samples were suspended in 15 µl of UHPLC loading buffer (3% acetonitrile + 0.1% trifluoroacetic acid).

Chromatographic separation was performed on a Dionex U3000 UHPLC system (Thermo Fisher Scientific, Darmstadt, Germany) equipped with an Acclaim PepMap 100 column (2 μm particle size, 75 μm × 500 mm) and µ-precolumn (300 μm × 5 mm) coupled online to Fusion Lumos mass spectrometer (Thermo Fisher scientific). The eluents used were; eluent A: 0.05% formic acid (FA), eluent B: 80% ACN + 0.05% FA. The separation was performed over a programmed 60-minute run. Initial chromatographic conditions were 4% B for 3 minutes followed by linear gradients from 4% to 50% B over 30 minutes, a 1-minute increase to 90% B, and 10 minutes at 90% B. Following this, an inter-run equilibration of the column was achieved by 15 minutes at 4% B. A constant flow rate of300 nl/min was used and 1 μl of sample was injected per run. Data acquisition on the Fusion Lumos mass spectrometer utilized HCD activation (NCE 30). A full scan MS acquisition was performed (resolution 120,000) scan range 300-1500 m/z, AGC target 4e5, maximum IT 50 ms. Subsequent data dependent MS/MS (resolution 30,000, minimum intensity 5e4, maximum IT 100 ms), of the most intense ions for the subsequent 3 seconds; MIPS mode was enabled for peptides, single charged and >7+ charged peptides were excluded, dynamic exclusion was enabled (60 sec duration); internal lock mass was enabled on 445.12003 m/z.

MS data files were searched against a combined set of fasta databases containing the full *M. mazei* proteome plus all predicted sORF encoded proteins (PTK_92x_Mmazei_Full_Plus_SEP_190425.fasta), and the Histag protein sequences provided for sp26 and GlnA (up26HisTagged+GlnAHis.fasta). In addition, protein sequences for the expression system, *E. coli* D3 and the cRAP list of common laboratory contaminants were included for all searches (PTK_proteome_Ecoli.fasta and cRAP47.fasta). The searches were performed using the Proteome Discoverer software package (version 2.2.0.388) using the SequestHT search algorithm. A semi-tryptic search was performed, (Precursor tolerance 10 ppm, fragment tolerance 0.02 Da, missed cleavages 2). Variable modifications; methionine oxidation and protein N-terminal acetylation, fixed modification; cysteine carbamidomethylation.

Strict parsimony criteria have been applied with a target FDR of 0.01 (1%) applied at the PSM, peptide, and protein level. In addition, proteins required two high confident peptides to be considered identified, or in the case of sP26, single peptides were manually validated to assure quality of the identification.

**Table S1:**
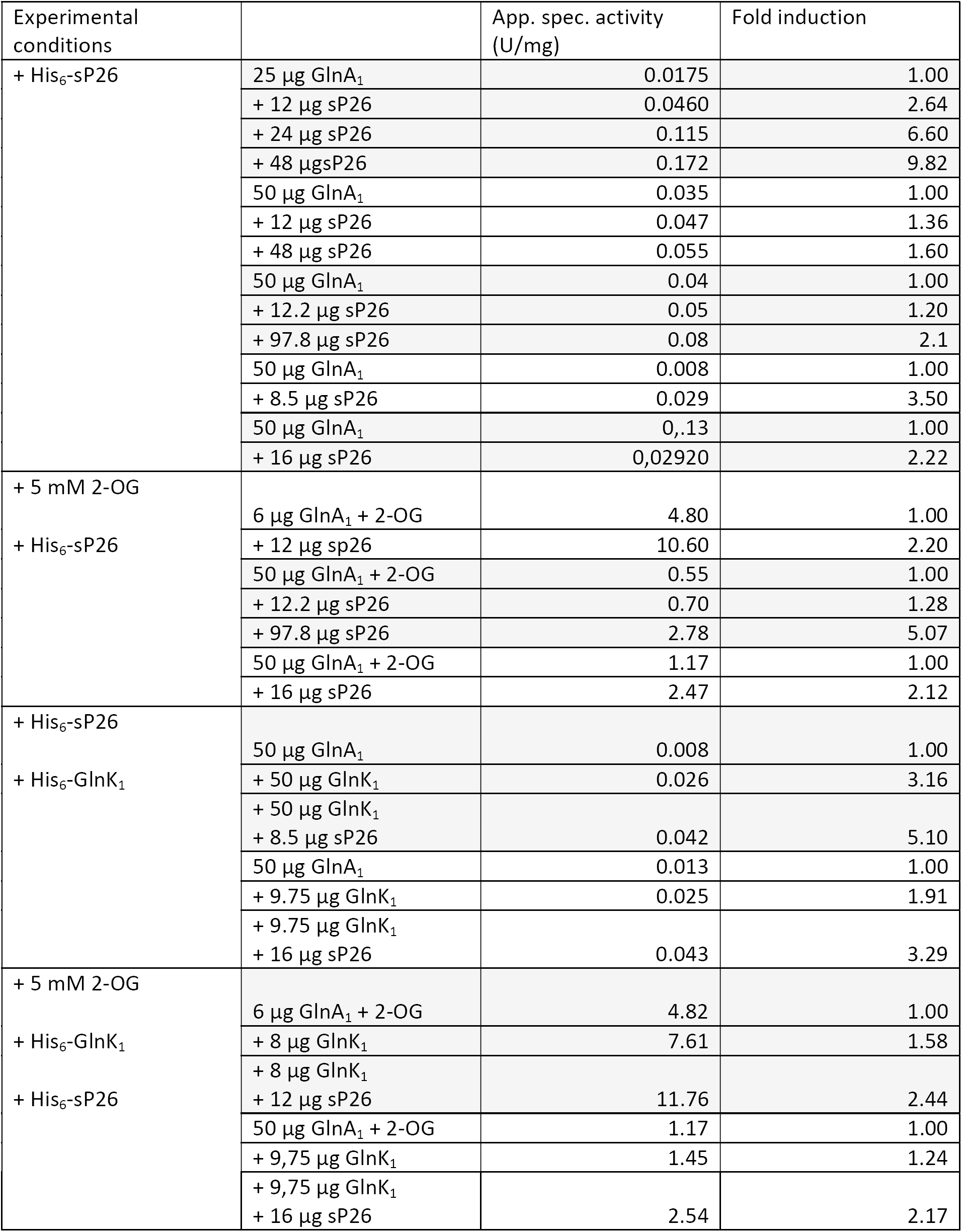
GlnA_1_ activity determined as described in Materials and Methods using purified proteins of independent purifications

| Experimental conditions |  | App. spec. activity (U/mg) | Fold induction |
| --- | --- | --- | --- |
| + His<sub>6</sub>-sP26 | 25 µg GlnA<sub>1</sub> | 0.0175 | 1.00 |
|  | + 12 µg sP26 | 0.0460 | 2.64 |
|  | + 24 µg sP26 | 0.115 | 6.60 |
|  | + 48 µg sP26 | 0.172 | 9.82 |
|  | 50 µg GlnA<sub>1</sub> | 0.035 | 1.00 |
|  | + 12 µg sP26 | 0.047 | 1.36 |
|  | + 48 µg sP26 | 0.055 | 1.60 |
|  | 50 µg GlnA<sub>1</sub> | 0.04 | 1.00 |
|  | + 12.2 µg sP26 | 0.05 | 1.20 |
|  | + 97.8 µg sP26 | 0.08 | 2.1 |
|  | 50 µg GlnA<sub>1</sub> | 0.008 | 1.00 |
|  | + 8.5 µg sP26 | 0.029 | 3.50 |
|  | 50 µg GlnA<sub>1</sub> | 0.13 | 1.00 |
|  | + 16 µg sP26 | 0.02920 | 2.22 |
| + 5 mM 2-OG<br>+ His<sub>6</sub>-sP26 | 6 µg GlnA<sub>1</sub> + 2-OG | 4.80 | 1.00 |
|  | + 12 µg sP26 | 10.60 | 2.20 |
|  | 50 µg GlnA<sub>1</sub> + 2-OG | 0.55 | 1.00 |
|  | + 12.2 µg sP26 | 0.70 | 1.28 |
|  | + 97.8 µg sP26 | 2.78 | 5.07 |
|  | 50 µg GlnA<sub>1</sub> + 2-OG | 1.17 | 1.00 |
|  | + 16 µg sP26 | 2.47 | 2.12 |
| + His<sub>6</sub>-sP26<br>+ His<sub>6</sub>-GlnK<sub>1</sub> | 50 µg GlnA<sub>1</sub> | 0.008 | 1.00 |
|  | + 50 µg GlnK<sub>1</sub> | 0.026 | 3.16 |
|  | + 50 µg GlnK<sub>1</sub><br>+ 8.5 µg sP26 | 0.042 | 5.10 |
|  | 50 µg GlnA<sub>1</sub> | 0.013 | 1.00 |
|  | + 9.75 µg GlnK<sub>1</sub> | 0.025 | 1.91 |
|  | + 9.75 µg GlnK<sub>1</sub><br>+ 16 µg sP26 | 0.043 | 3.29 |
| + 5 mM 2-OG<br>+ His<sub>6</sub>-GlnK<sub>1</sub><br>+ His<sub>6</sub>-sP26 | 6 µg GlnA<sub>1</sub> + 2-OG | 4.82 | 1.00 |
|  | + 8 µg GlnK<sub>1</sub> | 7.61 | 1.58 |
|  | + 8 µg GlnK<sub>1</sub><br>+ 12 µg sP26 | 11.76 | 2.44 |
|  | 50 µg GlnA<sub>1</sub> + 2-OG | 1.17 | 1.00 |
|  | + 9.75 µg GlnK<sub>1</sub> | 1.45 | 1.24 |
|  | + 9.75 µg GlnK<sub>1</sub><br>+ 16 µg sP26 | 2.54 | 2.17 |

